# Specialized parallel pathways for adaptive control of visual object pursuit

**DOI:** 10.1101/2025.04.23.650240

**Authors:** Matthew F. Collie, Chennan Jin, Victoria Rockwell, Emily Kellogg, Quinn X. Vanderbeck, Alexandra K. Hartman, Stephen L. Holtz, Rachel I. Wilson

## Abstract

To pursue a moving visual object, the brain must generate motor commands that continuously steer the object to the center of the visual field via feedback. The gain of this control loop is flexible, yet the biological mechanisms underlying such adaptive control are not well-understood. Here we show that adaptive control in the *Drosophila* pursuit system involves two parallel pathways. One detects objects in the periphery and steers them toward the center of the visual field. The other detects objects near the center of the visual field and steers them to the visual midline. This latter pathway is flexible: gain increases when the object is moving away from the midline and when the pursuer is running fast. This latter pathway is also preferentially recruited when the fly is aroused, and suppressing it decreases pursuit performance. Our findings demonstrate how adaptive control can emerge from parallel pathways with specialized properties.

## Introduction

A feedback control system works by continuously comparing its output to some setpoint, and then updating that output to correct any discrepancy, or “error”. For a given error, the size of this corrective change depends on the system’s gain, where higher gain produces more correction. Gain can be difficult to optimize: if it is too low, the system under-reacts to large errors, leading to sluggish performance, but if the gain is too high, the system over-reacts to small errors, resulting in overshooting and instability^1^. Moreover, the best compromise between these competing constraints can change over time.

A general solution to this problem is adaptive control, where the parameters of a feedback system are continuously adjusted to match current conditions^2^. With adaptive control, gain can be increased when rapid error correction is needed and reduced when stability is more important. Many engineered systems implement this through “gain scheduling”, assigning different gains for different conditions. Similarly, animals can dynamically adjust the gain of their visuo-motor feedback loop during visual pursuit—tracking or following a moving target—depending on their current behavioral state^3^. For example, in primates pursuing a target with their eyes, increased attention to the task causes an increase in pursuit gain^4–11^. Social interactions can drive particularly strong changes in attention or arousal; for instance, flying dragonflies increase visual pursuit gain when chasing rivals in territorial displays of physical prowess^12–14^.

Beyond gain scheduling, engineered feedback systems can also adapt by switching between different control strategies^2^. This, too, occurs in animal behavior: houseflies switch between two feedback control strategies when pursuing rivals^15^. In “direct pursuit”, the fly steers toward the rival’s current position, whereas in “anticipatory pursuit”, steering depends on both the rival’s position and his velocity on the pursuer’s retina, directing the fly toward the rival’s future location. Houseflies switch from direct pursuit to anticipatory pursuit when the rival moves into the central zone of their visual field^15^. This finding suggests that the central and peripheral zones are covered by different cell types, with different input sensitivities. Indeed, in dragonflies, pursuit is mediated by a set of descending neurons with diverse visual receptive fields^16–18^.

In short, flexible control is a widespread feature of visual object pursuit, and yet we do not fully understand the underlying neural mechanisms that generate flexible pursuit behavior. In this study, we investigate this problem by leveraging the *Drosophila melanogaster* connectomes^19–24^ and genetic toolbox. *Drosophila* pursue visual objects while flying or walking^25^, and the vigor of pursuit depends on behavioral state. For instance, appetitive odor can drive steering toward a visual object^26,27^, while food deprivation^28–31^ or social arousal^32–40^ can drive vigorous and persistent steering toward other flies during aggressive displays as well as courtship displays (**Fig. 1A**). Moreover, in *Drosophila*, arousal (and visual pursuit) can be evoked by stimulating defined neurons^29,37,39,41–45^, providing an experimental entry-point for investigating the mechanisms of adaptive pursuit.

**Figure 1:**
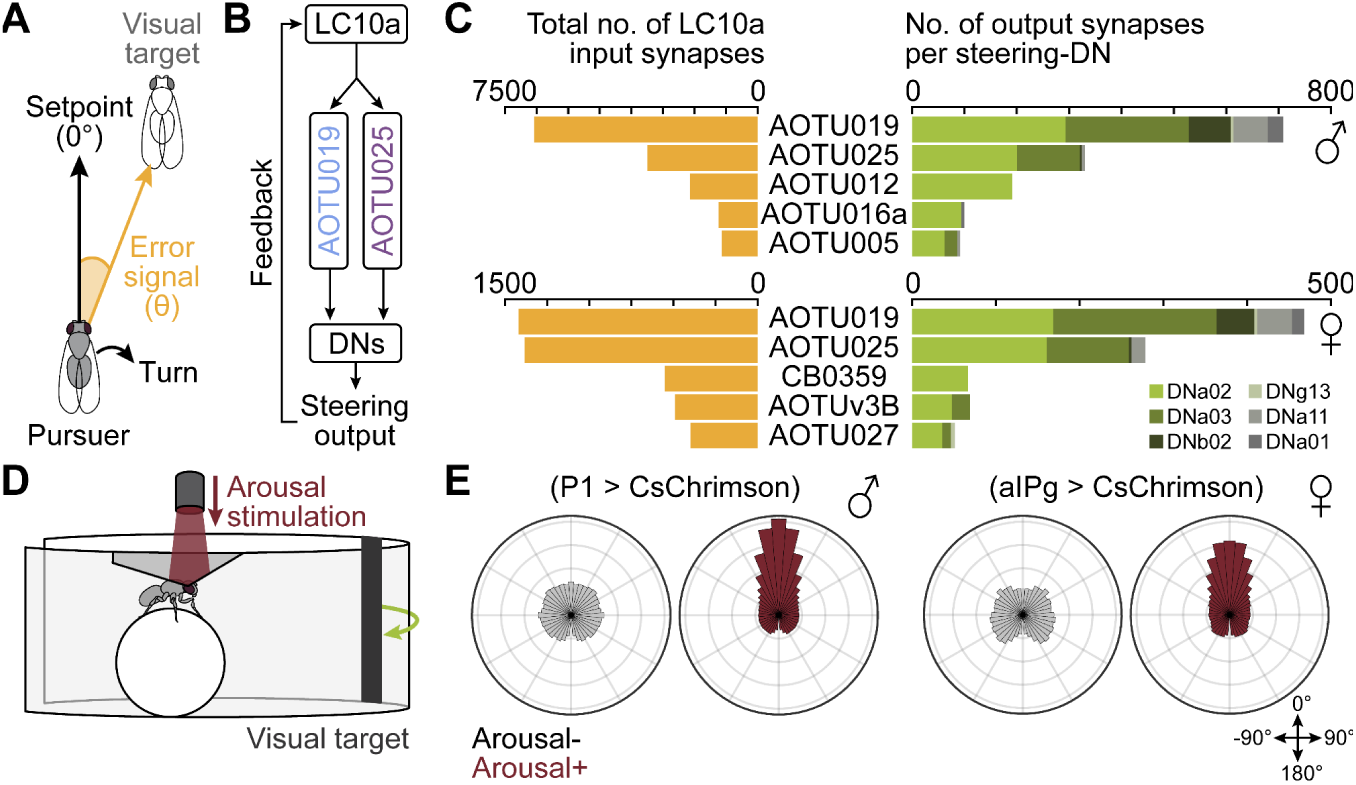
Shared circuitry for visual object pursuit in male and female *Drosophila*. **(A)** In visual pursuit, the pursuer’s task is to keep the target object (e.g., another fly) at the visual midline (0°). The angular deviation (θ) represents the error from this setpoint, which should drive corrective turning to re-center the object via feedback control. **(B)** Proposed feedback control model of visual pursuit in *Drosophila*. LC10a cells detect moving visual objects and influence steering descending neurons (DNs), largely via AOTU019 and AOTU025. **(C)** Top five AOTU cell types in males (top) and females (bottom) receiving direct LC10a input and providing direct outputs to walking-related steering DNs^48–50,57^. DNa03 primarily targets other DNs, including DNa02 (**Fig. S1B**). Synapse counts averaged across hemispheres. Connectivity in males (MaleCNS) and females (Flywire-FAFB). **(D)** Schematic of a fly walking on a spherical treadmill. In some experiments, we link the object’s position to the fly’s rotational velocity (“virtual reality” mode, i.e., “closed loop”). In others, we present moving objects independent of the fly’s rotational velocity (“open loop”). Arousal is evoked via optogenetic stimulation of P1 (males) or aIPg (females). **(E)** Polar histograms of object positions during virtual reality pursuit before and during arousal. Data are shown as probability densities (range: 0–0.085) and only include walking epochs (>1 mm/s). Arousal increased object concentration near the midline (smaller median absolute deviation from 0°) in both males (P1, n = 75, p < 0.0001) and females (aIPg, n = 21, p < 0.0001). Only males showed increased forward velocity (**Fig. S2A**).

Here we show how adaptive control of visual pursuit emerges from two parallel pathways. One of these pathways is engaged by a target object in the periphery, and it steers the object into the central region of the pursuer’s visual field. The other is engaged once the object enters the central region, and it steers the object to the visual midline. Notably, the latter pathway is preferentially recruited when the fly is aroused, and it increases steering in the direction of the target’s movement, while also increasing steering when the pursuer is running fast. Suppressing activity in this latter pathway decreases pursuit performance. Together, these results show how feedback systems can implement adaptive control via specialized parallel pathways whose gain depends on behavioral state.

## Results

### Two key AOTU cell types link object-directed steering with behavioral state

Visual signals for pursuit enter the central brain via LC10a cells, which detect small moving visual objects^40,45–47^. LC10a cells synapse onto neurons in the anterior optic tubercle (AOTU) that project to descending neurons that influence steering during walking^48–50^. Two AOTU cell types make the largest contribution to this pathway, namely AOTU019 and AOTU025 (**Fig. 1B,C, Fig. S1A,B**), but nothing is known about their physiology. The connectivity of AOTU019 and AOTU25 is largely similar in females and males (**Fig. 1C**), suggesting that these neurons represent core elements of the visual pursuit circuit in both sexes. Although pursuit behavior has sexually dimorphic features^29,32,51–53^, and it can be triggered by sex-specific neurons such as P1 (in males^37,39,41–45^) and aIPg (in females^29,53^), we find that basic visual pursuit is similar in males and females: P1 or aIPg stimulation drives steering toward a simple moving dark object in virtual reality (**Fig. 1D,E, Fig. S2A-E**). We use the term “arousal” to refer to the state of heightened visuo-motor engagement evoked by P1 or aIPg activation.

To investigate how these AOTU neurons contribute to pursuit behavior, and how their activity is modulated by arousal, we generated selective split-Gal4 lines for AOTU019 and AOTU025 (**Fig. S3A**) and targeted each cell type for electrophysiological recording while stimulating P1 or aIPg cells to evoke arousal. In both males and females, we found that arousal produced a large depolarization and increase in firing rate in AOTU019, but not in AOTU025 (**Fig. 2A,B**). Notably, aIPg cells make strong direct connections onto AOTU019^29^, but no connections onto AOTU025 (**Fig. S1C**). Similarly, P1 cells are positioned to exert strong disynaptic influence over AOTU019, but not AOTU025 (**Fig. S1D**). All these results indicate that arousal preferentially recruits AOTU019, suggesting it has a specialized function.

**Figure 2:**
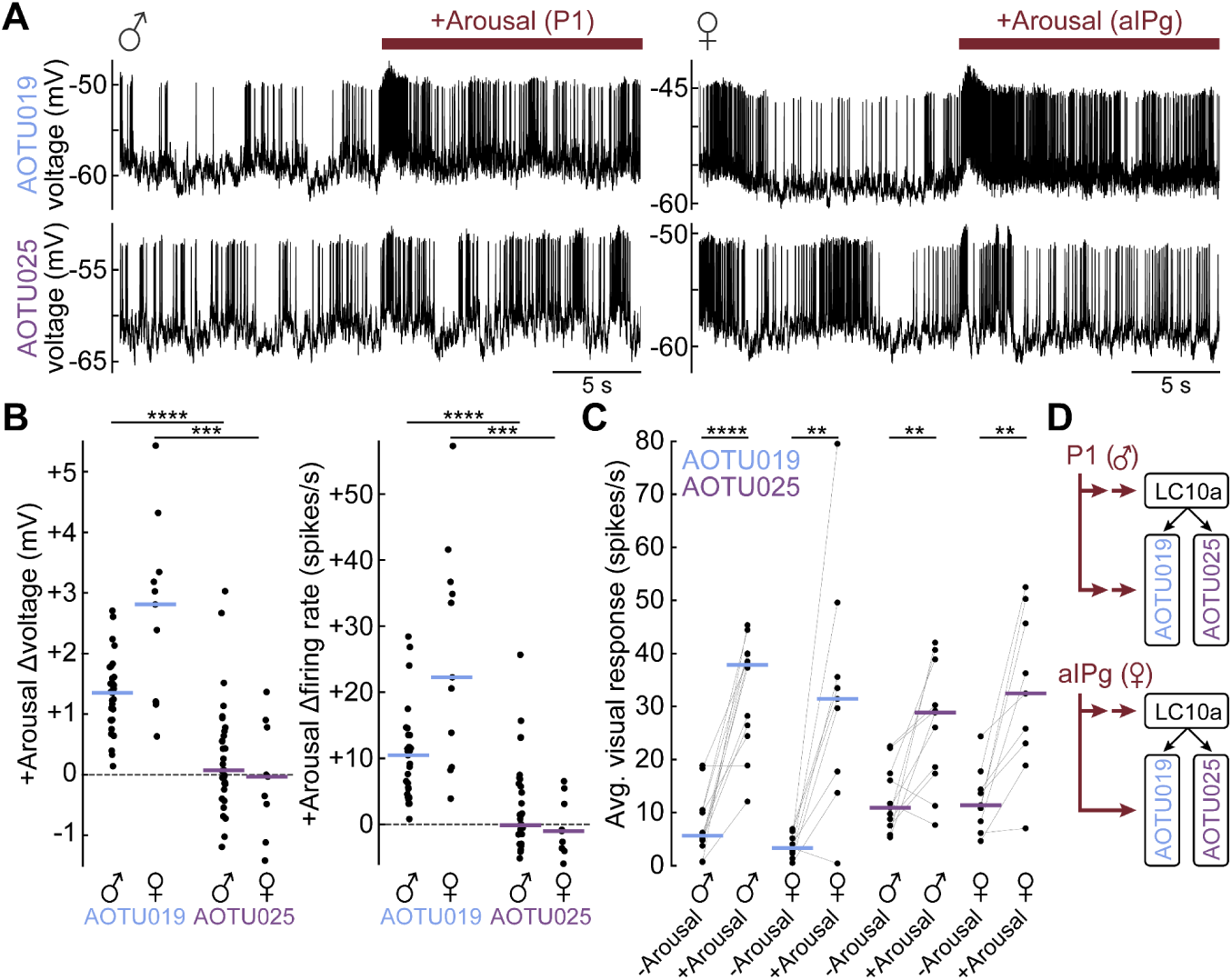
Arousal state differentially recruits AOTU visuo-motor pathways. **(A)** Example recordings from AOTU019 and AOTU025 cells before and during arousal stimulation. Effects were similar in males (left: P1) and females (right: aIPg). Right hemisphere recordings were made while flies walked in darkness. **(B)** Arousal-induced changes in membrane voltage and firing rate for AOTU019 and AOTU025 in males (P1: n = 31, 34) and females (aIPg: n = 11, 9). AOTU019 showed larger increases than AOTU025 (P1: p < 0.0001; aIPg: p < 0.001). Here and in **(C)**, activity was measured when flies were stationary and excluding the 250 ms after arousal onset/offset. Horizontal lines indicate medians. **(C)** Arousal-induced changes in visual responses for AOTU019 and AOTU025 in males (P1: n = 12, 11) and females (aIPg: n = 9, 9). Arousal increased visual responses (P1: p < 0.0001; aIPg: p < 0.001) with no significant difference between cell types (P1: p = 0.080; aIPg: p = 0.989). **(D)** In males, P1 cells disynaptically excite AOTU019, but not AOTU025 (**Fig. S1D**). In females, aIPg cells directly excite AOTU019, but not AOTU025 (**Fig. S1C**). In both sexes, arousal cells also gate LC10a visual responses^45,53^.

Next, we presented flies with moving visual objects (**Fig. S2F,G**), as these stimuli evoke strong responses in upstream LC10a cells^40,45–47^. We found these stimuli evoked almost no response in AOTU019 or AOTU025 without arousal, but with arousal, they evoked strong responses in both AOTU cell types (**Fig. 2C**). This is to be expected, as arousal is known to gate visual responses in LC10a cells^45,53^ (**Fig. 2D**).

Together, these results suggest that AOTU019 and AOTU025 have a central role in visual pursuit, transforming visual signals into descending steering drives. However, the activity of these cells depends on behavioral state, particularly for AOTU019 (**Fig. 2D**). These findings motivated us to investigate how exactly these AOTU cells transform visual signals into steering drives, and how they contribute to adaptive control.

### AOTU circuits divide visual space into parallel pathways

LC10a projections to the AOTU are retinotopic^46^, and each postsynaptic cell in the AOTU receives input from only a subset of the LC10a population^45^. In particular, we noticed that AOTU019 samples from LC10a cells devoted to the medial visual field, whereas AOTU025 samples from LC10a cells devoted to the lateral visual field (**Fig. 3A,B**). This anatomy suggests that AOTU019 detects objects near the center of the visual field, whereas AOTU025 detects objects in the periphery (**Fig. 3B**).

**Figure 3:**
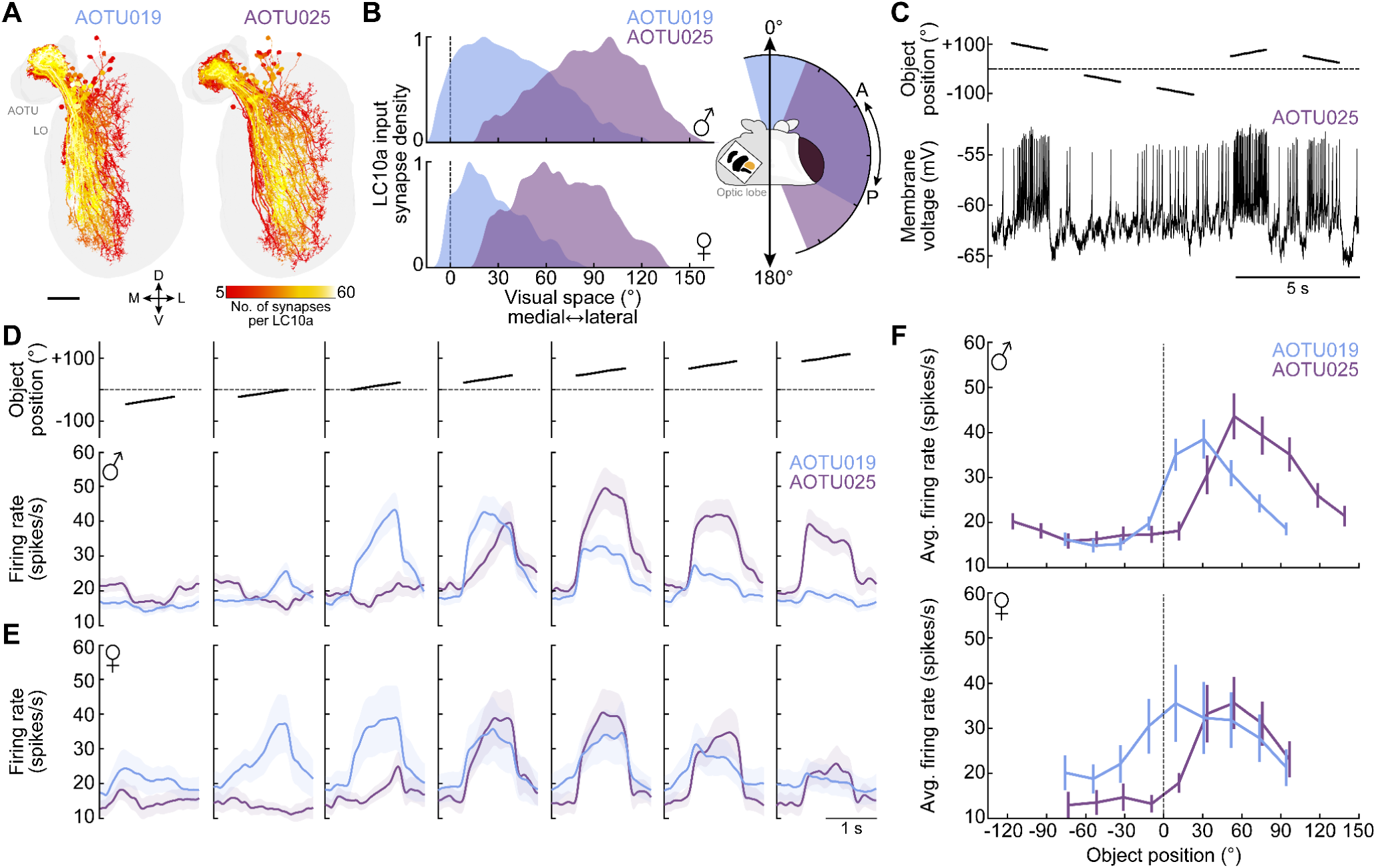
Parallel AOTU pathways encode partially overlapping regions of visual space. **(A)** LC10a cells with dendrites in the lobula (LO) and axons in the anterior optic tubercle (AOTU), color-coded by synapse count onto AOTU019 (left) or AOTU025 (right). Data from females (Flywire-FAFB). Scale bar: 20 µm. **(B)** Estimated visual receptive fields for AOTU019 and AOTU025 in males (top, MaleCNS) and females (bottom, Flywire-FAFB), derived from LC10a input density along the LO medial–lateral axis, including 15° binocular overlap^25,83^. Right: Estimated receptive fields (male) projected onto the anterior–posterior axis of the right retina. **(C)** Example AOTU025 recording during motion pulse presentations in a female with aIPg stimulation. Each pulse appeared at a random position and rotated 22.5° at 25 °/s before disappearing. Example male recordings in **Fig. S4A**. **(D)** Average motion pulse responses for AOTU019 (mean ± s.e.m., n = 25) and AOTU025 (n = 22) in males during P1 stimulation. Right-hemisphere cells were shown rightward pulses; left-hemisphere data were mirrored and combined. Response latencies: 40 ± 9 ms (AOTU019), 36 ± 10 ms (AOTU025). **(E)** Same as **(D)**, but for females during aIPg stimulation (AOTU019: n = 9, AOTU025: n = 9). Response latencies: 35 ± 7 ms (AOTU019), 52 ± 13 ms (AOTU025). **(F)** Average firing rate during motion pulses (mean ± s.e.m.) for AOTU019 and AOTU025 in males (top; n = 25, 22) and females (bottom; n = 9, 9). Object position corresponds to the pulse’s sweep center.

To test this idea, we made electrophysiological recordings from AOTU019 and AOTU025 and mapped their visual receptive fields by presenting flies moving objects on a cylindrical visual panorama (**Fig. 3C, Fig. S4A**). Specifically, objects appeared at randomized positions and then transiently moved left or right before disappearing; we refer to these brief visual stimuli as “motion pulses”. In these experiments (and in the rest of this study), we used arousal stimulation to ensure that AOTU neurons are visually responsive.

We found that AOTU019 responded best to motion pulse stimuli near the center of the visual field (**Fig. 3D-F**). Conversely, AOTU025 responded best to motion pulse stimuli in the periphery (**Fig. 3D-F**). We also found considerable overlap in the receptive fields of these cell types, meaning they will often be co-recruited. Notably, the visual receptive field properties of AOTU019 and AOTU025 were similar in males and females (**Fig. 3F**). In sum, these results suggest that moving visual objects evoke pursuit behavior via parallel visuo-motor pathways, which divide the visual field into overlapping central and peripheral zones.

### Steering can arise from asymmetric excitation or inhibition

Next, we asked how visual responses in AOTU neurons are translated into the steering commands during visual pursuit. Here, it is relevant that these two AOTU cell types release different neurotransmitters: AOTU025 is cholinergic, whereas AOTU019 is GABAergic. This conclusion is based on machine classification of electron microscopy (EM) images^54^, as well as genetic evidence^55^ (**Fig. S3B**). This would imply that AOTU025 is excitatory, whereas AOTU019 is inhibitory.

AOTU025 projects to the ipsilateral copies of steering descending neurons (DNs), whereas AOTU019 targets the contralateral copies of these DNs (**Fig. 4A, Fig. S5A**). Each type of steering DN is represented by a single pair of mirror-symmetric neurons—one per hemisphere^56^—and when their firing rates are unequal, the fly turns toward the side of higher DN activity^48,49,57–59^, with a rotational velocity that scales with the right-left firing rate difference^48,57^. Given their neurotransmitter profiles and their connection patterns, both AOTU cells are predicted to drive higher DN firing on the side of the visual object, and thus turning toward the object (**Fig. 4A**). For example, an object on the right will excite both AOTU025 and AOTU019 in the right hemisphere, leading to excitation of right DNs and inhibition of left DNs, respectively. Both effects should increase the right-left difference in DN activity, driving rightward turning. It might seem odd to think that inhibitory neurons can drive downstream motor programs, but there is good evidence for this notion in other systems^60^.

**Figure 4:**
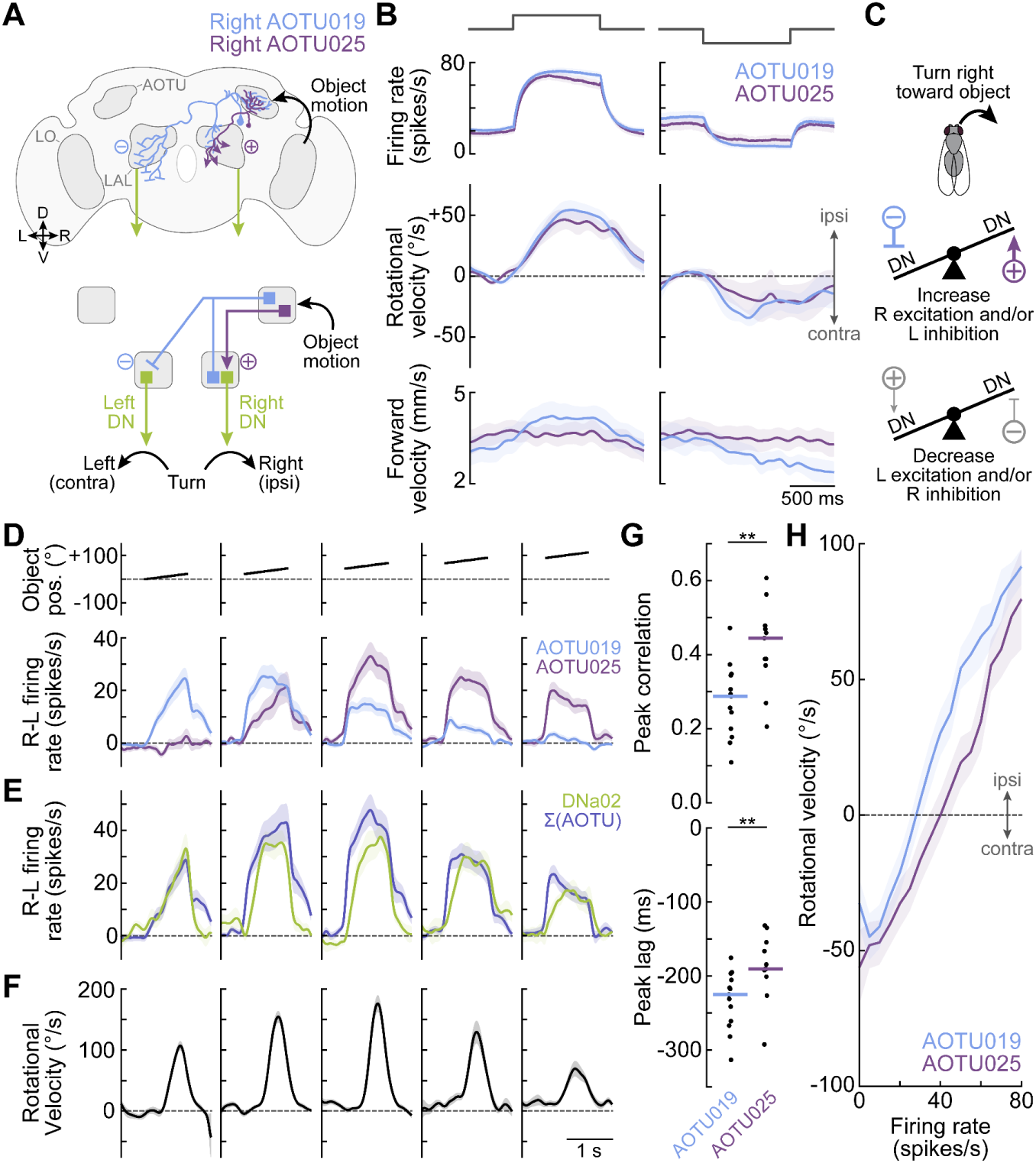
Steering can arise from asymmetric excitation or inhibition. **(A)** Front-view brain schematic (top) and circuit diagram (bottom). AOTU019 and AOTU025 link object motion to steering DNs (e.g., DNa02, **Fig. 1C**) in premotor regions (e.g., lateral accessory lobe; LAL). AOTU019 is inhibitory and projects contralaterally, with dendrites in both the AOTU and LAL; AOTU025 is excitatory and projects ipsilaterally, with dendrites in the AOTU only (**Fig. S5**). “Cell on the right” refers to soma and dendrites in the right hemisphere. **(B)** Effects of current injection into AOTU019 (+: n = 15; −: n = 8) or AOTU025 (+: n = 15; −: n = 5) during arousal stimulation and walking. Depolarizing either cell type drove ipsilateral turning (p < 0.0001), and for AOTU019 increased forward velocity (p < 0.005). Hyperpolarizing AOTU019 drove contralateral turning (p < 0.05) with a trend toward reduced forward velocity (p = 0.066). Hyperpolarizing AOTU025 had no significant effects (p = 0.362, p = 0.246). Ipsilateral indicates toward the recorded soma. **(C)** Schematic of “see-saw steering control” via AOTU inputs. Steering right can result from increasing excitation to the right, decreasing excitation to the left, increasing inhibition to the left, or decreasing inhibition to the right. If DNs linearly sum their inputs and steering depends on the right–left difference in DN activity, ipsilateral excitation and contralateral inhibition will have equivalent effects. **(D)** Blue: Motion pulse responses for AOTU019 (mean ± s.e.m., n = 23), shown as the right–left firing rate difference (R–L; see Methods). This reflects the net steering drive, or “tilt” in the see-saw in **(C)**, where positive values indicate rightward drive. Magenta: Same for AOTU025 (n = 22). Responses in **(D-H)** are recorded during arousal stimulation. **(E)** Green: R–L difference in DNa02 firing rate (mean ± s.e.m., n = 11). Purple: Sum of AOTU019 and AOTU025 responses from **(D)** (mean ± s.e.m. with error propagation), representing their combined steering drive. **(F)** Mean turn responses to rightward motion pulses (± s.e.m., n = 40). **(G)** Peak cross-correlation between firing rate and rotational velocity during motion pulses for AOTU019 (n = 13) and AOTU025 (n = 11). Both cells preceded turning, with AOTU025 showing stronger correlation (p < 0.01) and shorter lag (p < 0.01). Horizontal lines indicate medians. **(H)** Mean rotational velocity (± s.e.m.) binned by firing rate 200 ms prior, for AOTU019 (n = 13) and AOTU025 (n = 11). No significant difference between cell types (p = 0.156).

To test this prediction, we injected current via the recording electrode into individual AOTU cells, and measured the resulting change in the fly’s rotational velocity as the fly walked on a spherical treadmill. Here, and in the rest of this study, we focused on male flies, as AOTU019 and AOTU025 have similar properties in males and females, but males produce tighter fixation on the visual object (**Fig. 1E, Fig. S2A**). We found that direct excitation of either AOTU019 or AOTU025 was sufficient to produce ipsilateral turning (i.e., turning toward the side of the excited cell; **Fig. 4B**). Conversely, when we injected hyperpolarizing current to inhibit these cells, we observed contralateral turning (**Fig. 4B**). The average size of the behavioral response was similar for the two cell types.

Together, these results imply that the same motor command can arise through multiple mechanisms: for example, a rightward turn with a specific rotational velocity can result from increased excitatory drive to DNs on the right, decreased excitatory drive to DNs on the left, increased inhibitory drive to DNs on the left, decreased inhibitory drive to DNs on the right, or any combination of these. We call this the “see-saw model of steering control”^48^ (**Fig. 4C**). In visual circuits, this motif (i.e., “crossover inhibition”) improves robustness to noise^61^, and the same may be true in motor control circuits.

Unexpectedly, we also found that depolarization of AOTU019 increased forward velocity, whereas depolarization of AOTU025 had no effect on forward velocity (**Fig. 4B**). This cannot be explained as a direct consequence of faster turning in one group of flies, because the two groups had similar rotational velocity responses (**Fig. 4B**). Rather, these results imply that AOTU019 actually accelerates the pursuer’s forward movement toward a visual object, while also keeping the pursuer oriented toward that object. It makes sense that AOTU025 should not accelerate forward movement, because the pursuer should not accelerate until he has successfully oriented toward the object.

Importantly, we can accurately predict the motion pulse responses of steering DNs and the fly’s rotational velocity, based on the activity of these two AOTU cells. Here we focused on DNa02, because this DN strongly correlates with (and influences) steering in walking flies^48,49,57^, and it has strong direct and indirect input from AOTU019 and AOTU025 (**Fig. 1C, Fig. S1A,B**). When we used electrophysiological recordings to measure DNa02 activity during arousal stimulation, we found that the combined visually-evoked activity of the four AOTU neurons (AOTU019 and AOTU025, in the right and left hemispheres) could predict the visually-evoked right-left asymmetry in DNa02 activity (**Fig. 4D,E**). We also found that visual responses in DNa02 could predict the fly’s rotational velocity in response to motion pulse stimuli (**Fig. 4F, Fig. S6**). While steering is likely influenced by multiple DNs^48–50,57,58^, DNa02 can still be predictive of steering behavior if its visual responses are correlated with those of other co-activated steering DNs. Notably, AOTU019 and AOTU025 firing rate changes typically precede changes in the fly’s rotational velocity (**Fig. 4G**), and they correlate with rotational velocity in a graded manner (**Fig. 4H**).

Taken together, our results thus far imply a clear division of labor in this circuit. AOTU025 detects an object in the periphery of the visual field, and it steers the object into the central zone. Next, AOTU019 detects an object inside the central zone, and it steers the object to the midline, while also increasing the pursuer’s forward velocity.

### Direction selectivity is highest around the center of the visual field

In principle, pursuit performance would be better if the brain took account of the object’s direction of movement. Indeed, we confirmed that visual pursuit behavior is direction-selective, as reported previously^45^: flies turned more vigorously toward a motion pulse if it was moving away from the midline (progressive motion), as compared to a pulse in the same position but moving toward the midline (regressive motion; **Fig. 5A**). We found that this behavioral bias is stronger for motion pulses in the central zone of the visual field, as compared to more peripheral motion pulses. Notably, the central zone is the region where AOTU019 cells are most active (**Fig. 3F**). Indeed, we found that AOTU019 cells are direction-selective and respond best to progressive object motion (**Fig. 5B**). As a result, the interhemispheric difference in AOTU019 firing rate is larger when an object is moving away from the midline, as compared to an object moving toward the midline (**Fig. 5B,C**). By comparison, we found that AOTU025 cells are not direction-selective (**Fig. 5B,C**).

**Figure 5:**
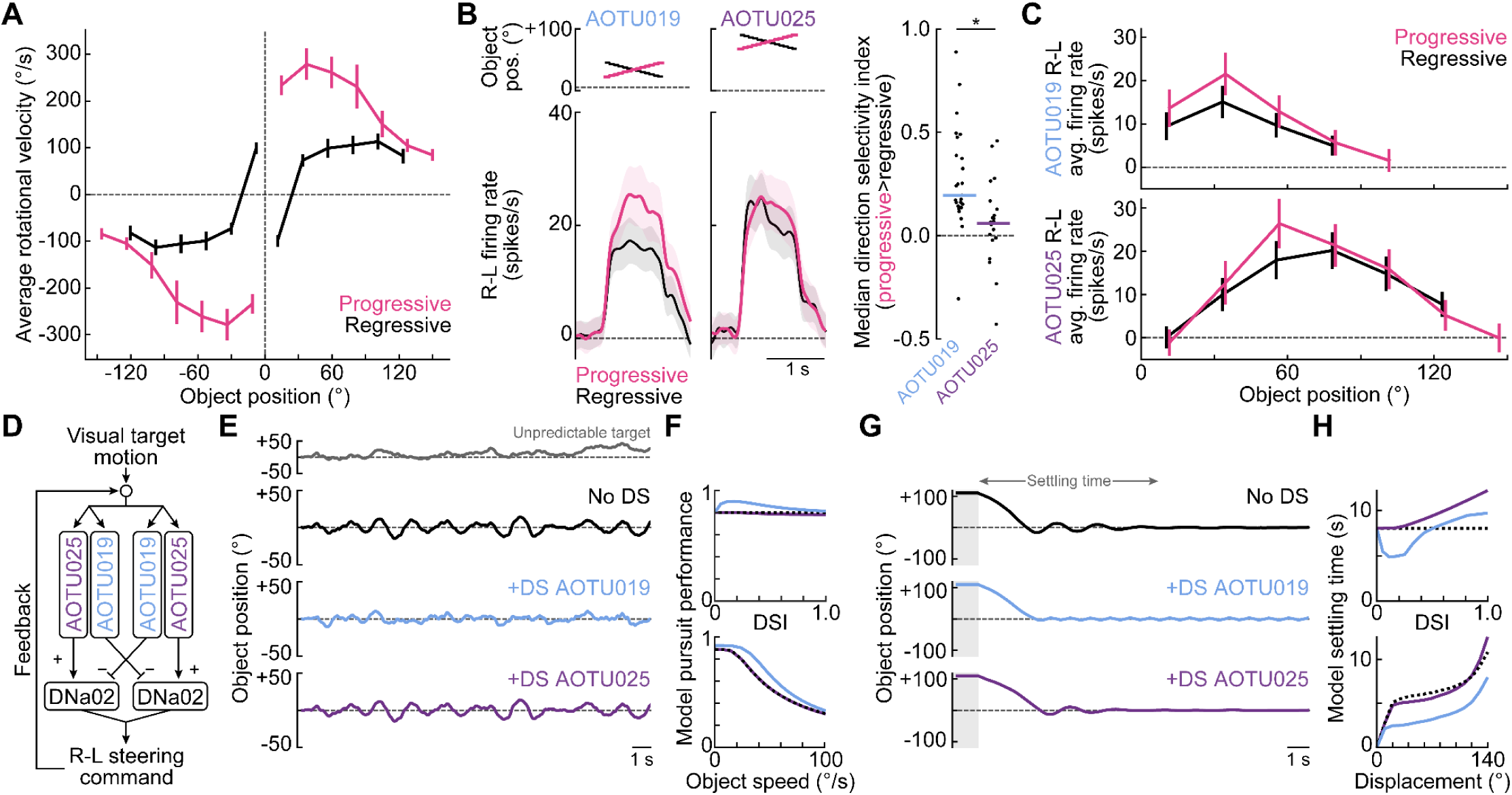
Direction selectivity is highest around the frontal visual field. **(A)** Behavioral responses progressive versus regressive motion pulses (mean ± s.e.m., n = 16 flies). The effect of motion direction depends significantly on object position (p < 0.0001). Motion pulse speed: 25 °/s. Data from intact (head-closed), arousal-stimulated males. Responses to left and right stimuli were mirrored and averaged. **(B)** R-L firing rate differences evoked by progressive versus regressive motion pulses at the preferred position for AOTU019 (mean ± s.e.m., n = 25) and AOTU025 (n = 22). Right: Median direction selectivity index (DSI) per fly, averaged across object positions that elicit a neural response. Positive DSI indicates progressive motion preference. DSI is higher for AOTU019 versus AOTU025 (p < 0.05). Horizontal lines indicate medians. **(C)** R-L firing rate differences across all object positions in the right hemifield for AOTU019 (n = 25) and AOTU025 (n = 22). AOTU019 responses varied with motion direction (p < 0.05), whereas AOTU025 responses did not (p = 0.667). Both cell types showed a trend toward greater direction selectivity in the central visual field (interaction: p = 0.190, p = 0.081). **(D)** Model of pursuit behavior: Connectome data specify AOTU019 and AOTU025 receptive fields and their output weights onto DNa02 (Fig. 1C). The R-L difference in DNa02 activity drives steering 200 ms later (Fig. 4G), which updates the object’s position and drives changes in AOTU activity. At each time step, the object was configured to either move unpredictably along a random trajectory **(E-F)** or hold at a fixed position before releasing **(G-H)**. **(E)** Object position in the pursuer’s visual field for three model variants: no direction selectivity (DS), DS in AOTU019 only, or DS in AOTU025 only. In all cases, feedback control steers the object toward the midline (0°). However, DS in AOTU019 maintains tighter fixation. The gray trace shows the object trajectory for an immobile pursuer (example mean object speed = 25 °/s). **(F)** Model performance for tracking an unpredictable target. DS in AOTU019 improves performance for small values of DSI (top, for object speed = 25 °/s) across object speeds (bottom, DSI = 0.2). Dashed line indicates no DS. Performance is quantified as the probability of the object remaining within ±5° of the midline. **(G)** Same as **(E)** but with the object initially displaced and held for 1 s (gray shading) before release. **(H)** Model performance for tracking displaced objects. DS in AOTU019 decreases settling time for small values of DSI (top, for object displacement = 120°)across different displacements (bottom, DSI = 0.2). Settling time is defined as the time to remain within ±2° for >1 s.

To show why direction selectivity is particularly useful around the center, we can use our network model (**Fig. 4A**) to simulate visual pursuit behavior (**Fig. 5D**). In this simulation, the pursuer’s orientation toward the object updates over time while maintaining a fixed distance. At each time point, the object’s position on the pursuer’s retina drives responses in two visual pathways, one central and one peripheral, based on our data (**Fig. 1C, 3B**). The combined activity of these pathways generates the pursuer’s predicted steering signal, which updates the object’s position on the retina via feedback.

As one might expect, our simulations show that this feedback loop brings the object toward the visual midline, even if the visual system is not direction-selective (**Fig. 5E-H**). Adding direction selectivity to the central pathway improves pursuit performance: the object is maintained closer to the center (**Fig. 5F**), and the system restabilizes more quickly after abrupt object movements (**Fig. 5H**). By contrast, adding direction selectivity to the peripheral pathway offered no performance benefit (**Fig. 5F**) and it even delayed the corrective response after larger displacements from the midline (**Fig. 5H**). This is because a large error demands a rapid corrective turn regardless of the object’s direction of motion.

### Faster running increases steering and visual responses near the visual field center

During visual pursuit, flies vary not only their steering, but also their forward velocity^35,40,62^. Indeed, in virtual reality experiments, we found that epochs of more consistent orienting toward the object coincided with epochs of faster forward velocity (**Fig. 6A,B**). Moreover, a given visual object position evoked stronger steering responses when forward velocity was also high (**Fig. 6C,D**).

**Figure 6:**
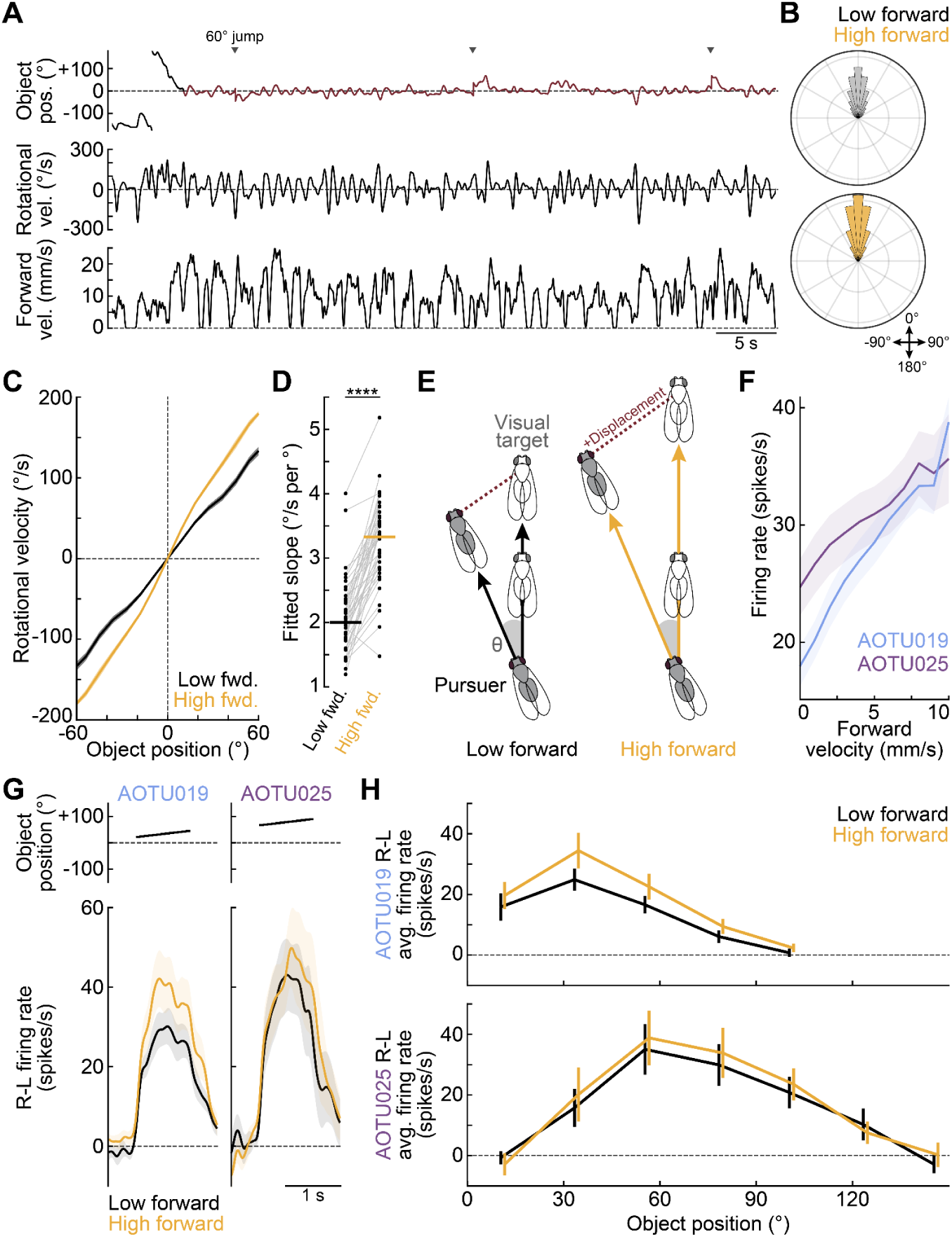
Increasing forward velocity increases steering gain. **(A)** Example of an arousal-stimulated male pursuing a visual object in virtual reality. Red marks continuous pursuit epoch. Triangles denote ±60° object “jumps” used to sample a wider range of object positions and evoke compensatory turns. **(B)** Polar histogram of visual object positions during pursuit for low (bottom third) and high (top third) forward velocity epochs (n = 40). Data shown as probability densities (range: 0–0.175). At higher forward speeds, object positions were more concentrated near the midline (reduced median absolute deviation from 0°: p < 0.0001). **(C)** Rotational velocity during pursuit, binned by object position in the frontal field (<60°), for low and high forward velocity epochs (mean ± s.e.m., n = 40). Responses to left and right stimuli were mirrored and averaged. **(D)** Steering gain, defined as the slope of the relationship between rotational velocity and object position, was significantly higher at faster forward velocities (n = 40, p < 0.0001). Horizontal lines indicate medians. **(E)** Higher forward velocity increases the impact of a given angular error (θ) on the pursuer’s path. Here, the pursuer and target fly are schematized as having the same forward velocity. **(F)** Firing rate binned by forward velocity during motion pulse presentation for AOTU019 (mean ± s.e.m., n = 25) and AOTU025 (n = 20). Cell types differed significantly (p < 0.05), with spiking varying by velocity in a cell type–specific manner (interaction, p < 0.0001). Changes in AOTU019 firing were typically synchronous with forward velocity changes (median lag: –3.0 ± 8.5 ms), whereas AOTU025 showed a more binary relationship to movement. **(G)** Visual responses to motion pulses at each cell’s preferred object position during low (<median) and high (>median) forward velocity epochs for AOTU019 (mean ± s.e.m., n = 15) and AOTU025 (n = 8). **(H)** R-L visual response means (± s.e.m.) to motion pulses during low and high forward velocity epochs for AOTU019 (n = 15) and AOTU025 (n = 8). Higher forward velocity significantly increased the R-L response in AOTU019 (p < 0.01), but not AOTU025 (p = 0.768).

This relationship makes sense: increased steering and increased forward velocity both bring the pursuer closer to his target. Moreover, vigorous steering becomes more important at higher forward velocity, because any angular error produces a faster-growing deviation in the pursuer’s path (**Fig. 6E**). For both these reasons, it is logical that increases in forward velocity are associated with increased steering gain.

In addition, we found that spontaneous increases in forward velocity are associated with increases in AOTU responses to motion pulse stimuli. This effect was graded and nearly linear (**Fig. 6F**), and it was significantly stronger for AOTU019, as compared to AOTU025. Moreover, during faster running, motion pulses produced larger interhemispheric differences in firing rate for AOTU019, but not AOTU025 (**Fig. 6G,H**).

To summarize, increased forward velocity is correlated with increased steering gain, stronger visual responses in AOTU019, and increased interhemispheric difference in AOTU019 activity. AOTU019 firing rate fluctuations are essentially synchronous with changes in forward velocity, suggesting that AOTU019 is receiving a copy of forward velocity commands, rather than always driving these commands. In this regard, it is notable that AOTU019 has a large dendritic arbor in the lateral accessory lobe (LAL; a premotor brain region) whereas AOTU025 does not (**Fig. 4A**, **Fig. S5A-C**). That said, direct depolarization of AOTU019 (but not AOTU025) can also drive increases in forward velocity (**Fig. 4B**). This is logical: the pursuer should increase their forward velocity only if they are successfully orienting toward their target, but not if the target remains in their visual periphery. Taken together, all these data imply a positive feedback loop linking steering with forward velocity. When the pursuer succeeds in orienting toward their target, this triggers forward acceleration, which then increases the vigor of ongoing corrective steering commands, while also sustaining ongoing fast running.

### Suppressing the central visual pathway reduces pursuit

All our results thus far point to a specialized role for AOTU019 in visual pursuit. We therefore asked whether suppressing AOTU019 interferes with pursuit behavior. We expressed the inwardly rectifying potassium channel Kir2.1 in AOTU019 cells (**Fig. S7A,B**), and we compared these flies to genetic controls bearing only the UAS-Kir2.1 transgene or the AOTU019 split-Gal4 line. In virtual reality experiments, suppressing AOTU019 activity produced a significant decrease in the fly’s rotational velocity in the direction of the object (**Fig. 7A,B**). This effect was small, consistent with the predictions of our network model (**Fig. 7C**), provided that this model includes several minor AOTU cell types whose visual receptive fields are partially overlapping with that of AOTU019 (**Fig. 1C, Fig. S8**).

**Figure 7:**
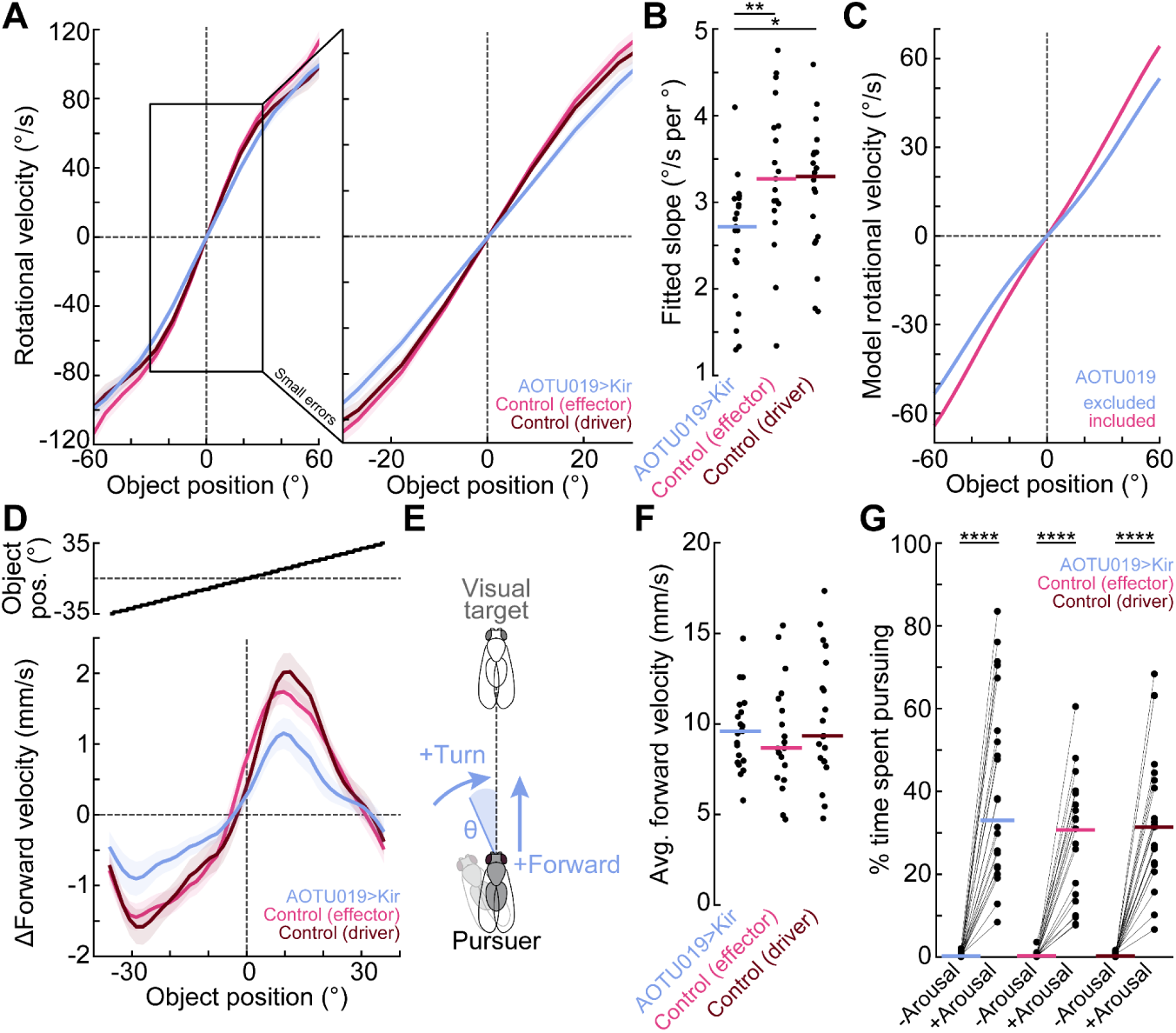
Silencing the frontal pathway impairs object-directed steering during arousal. **(A)** Rotational velocity binned by object position in the frontal field (<60°) during virtual reality pursuit for AOTU019>Kir (n = 21), effector control (n = 19), and driver control (n = 21) flies (mean ± s.e.m.). Right: Expanded view near the midline shows reduced steering toward the object in AOTU019>Kir flies. Responses to left and right stimuli were mirrored and averaged. **(B)** Steering gain (the slope of the relationship between rotational velocity and object position) was significantly reduced with AOTU019 silencing (p < 0.05, p < 0.01). Horizontal lines indicate medians. **(C)** Simulation summary showing rotational velocity versus object position, with AOTU019 either included or excluded. Excluding AOTU019 reduces steering toward the object. Minor LC10a-to-DNa02 pathways included (Fig. 1C**, Fig. S8**). **(D)** Change in forward velocity binned by object position for an open-loop stimulus for AOTU019>Kir, effector control, and driver control flies (mean ± s.e.m., n = 20 each). Changes are relative to each fly’s baseline. A significant genotype-position interaction (p < 0.0001) means that the effect of object position on forward velocity is significantly altered in AOTU019>Kir flies. This indicates that AOTU019 contributes to the normal sharp forward acceleration that occurs when an object sweeps across the midline. **(E)** Schematic illustrating how AOTU019 activity promotes object-directed steering and forward acceleration once oriented. **(F)** Distribution of average forward velocity during virtual reality pursuit for AOTU019>Kir (n = 21), effector control (n = 19), and driver control (n = 21) flies. No significant difference across genotypes (p = 0.477). **(G)** Percentage of time spent pursuing a visual object during virtual reality pursuit for AOTU019>Kir (n = 21), effector control (n = 19), and driver control (n = 21) flies before and during arousal stimulation. P1 stimulation increased pursuit in all genotypes (p < 0.0001, Bonferroni-corrected), with no effect of genotype (p = 0.999).

Recall that direct stimulation of AOTU019 drives not only object-directed steering, but also faster running (**Fig. 4B**). Given this, we might predict that AOTU019 activity is required for the fly to accelerate forward when oriented toward the visual object. To test this, we presented an object oscillating back and forth across the visual field. In control flies, we found a reliable forward acceleration as the object crossed the midline, but this response was significantly blunted by suppressing AOTU019 (**Fig. 7D**).

Together, these results argue that AOTU019 contributes to object-directed steering in the central visual field, as well as faster running when the visual object enters the central visual field (**Fig. 7E**). It is notable that suppressing AOTU019 has any phenotype, as there is only one AOTU019 cell in each hemisphere, and this is only one of 14 cells postsynaptic to LC10a and presynaptic to DNa02^45^ (**Fig. S8**). Lastly, we confirmed that these phenotypes were not attributable to a generalized locomotor defect: there was no effect of silencing AOTU019 on overall forward velocity (**Fig. 7F**), probability of pursuit engagement (**Fig. 7G**), or optomotor responses (**Fig. S7D**).

### AOTU pathways predict steering in darkness

Thus far, we have focused on neural and behavioral responses to visual stimuli. However, we found that even in darkness, firing rate changes in both AOTU019 and AOTU025 correlated with changes in the fly’s rotational velocity (**Fig. 8A,B**). In darkness, just as in our visual stimulus experiments, we found that arousal increased the correlation between firing rate and rotational velocity, and in the aroused state, this correlation was not significantly different from that measured during visual object pursuit (**Fig. 8B**). Moreover, in darkness, with arousal, the graded relationship between firing rate and rotational velocity was similar to what we observed during visual object pursuit (**Fig. 8C**).

**Figure 8:**
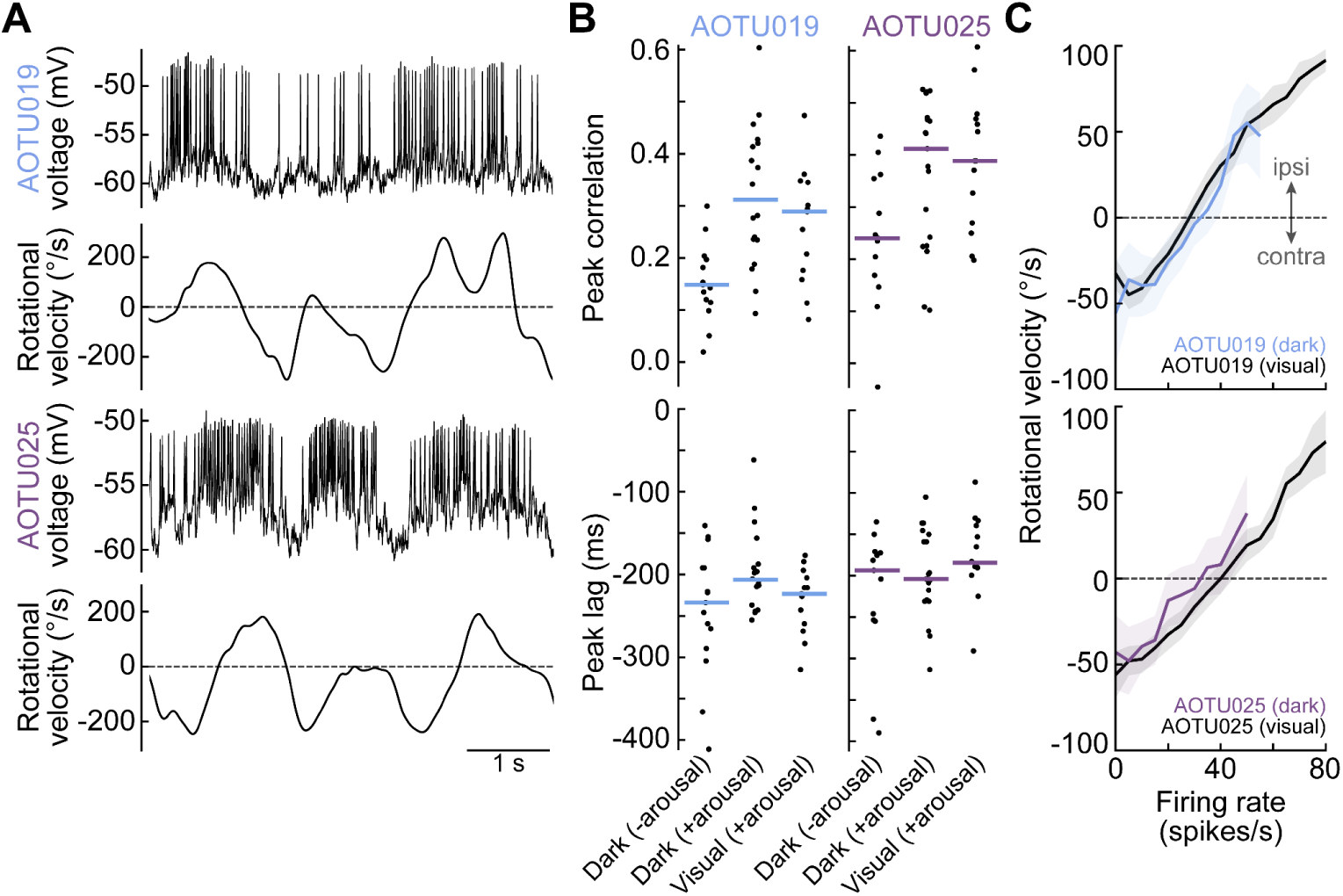
AOTU pathway activity correlates with steering in darkness. **(A)** Example recordings from AOTU019 and AOTU025 in darkness during arousal stimulation, showing increased firing preceding ipsilateral turns. Both cells were recorded on the right. **(B)** Peak cross-correlation between firing rate and rotational velocity in males walking in darkness (before or during arousal) or during motion pulse presentation (during arousal) for AOTU019 (n = 18, 13) and AOTU025 (n = 19, 14). Arousal significantly increased peak correlation for both AOTU019 (p < 0.0001) and AOTU025 (p < 0.001). Lag timing was modestly affected for AOTU019 (p < 0.01) but not for AOTU025 (p = 0.245). During arousal, peak cross-correlations did not differ between darkness and visual conditions for either cell type (AOTU019: correlation p = 0.141, lag p = 0.193; AOTU025: correlation p = 0.860, lag p = 0.285). Horizontal lines indicate medians. Data from visual stimulus conditions are also shown in Fig. 4G. **(C)** Mean rotational velocity (± s.e.m.) binned by firing rate 200 ms prior for AOTU019 (top: n = 12 in darkness, 13 during motion pulses) and AOTU025 (bottom: n = 11 in darkness, 11 during motion pulses). Data in black reproduced from Fig. 4H.

Importantly, in darkness, AOTU019 and AOTU025 activity preceded turning by about 200 ms, consistent with the idea that these AOTU neurons have a causal role in turning (**Fig. 8B**). Thus, it seems that these circuits receive non-visual inputs as well as visual inputs, and they likely contribute to steering in darkness as well as in the presence of a visual stimulus. The effect of visual stimuli is simply to evoke higher AOTU firing rates and larger rotational velocity fluctuations (**Fig. 8C**).

## Discussion

Visual pursuit (tracking or following a moving object) is a basic problem in visuo-motor control. In many species, behavioral studies have shown that visual pursuit involves adaptive control—the continuous adjustment of control system parameters to suit current conditions^3–15^. In *Drosophila*, visual pursuit behavior can be adaptively modulated by social arousal, hunger, and other cues^25–40^. Here we show how adaptive control in the *Drosophila* pursuit system emerges from two parallel pathways with specialized roles: one steers objects from the periphery toward the center, while the other maintains central object fixation. This organization enables flexible control gain and control logic. In particular, we describe mechanisms for adaptive control of pursuit based on four factors: (1) object position, (2) object direction of motion, (3) the pursuer’s running speed, and (4) the pursuer’s arousal state.

First, the position of the visual object determines which pathway is engaged: an object in the central visual field recruits AOTU019, whereas an object in the periphery recruits AOTU025. This architecture allows the visual pursuit system to respond differently to small versus large errors. Notably, AOTU019 and AOTU025 have overlapping visual receptive fields, so as an object moves from the periphery toward the midline (or vice versa), these AOTU neurons will be co-activated, ensuring a smooth handoff from one cell type to the other. In systems with gain scheduling, engineers use interpolation to create a smooth transition between different gains^2^. In effect, the network we describe here implements a graded interpolation between control parameters by ensuring co-activation at the boundary between the central and peripheral visual field.

A second key factor is the direction of object movement relative to the pursuer’s visual midline. Consistent with previous work^45^, we show that movement away from the midline drives a larger steering response than movement toward the midline. In effect, the pursuer anticipates the future position of the visual object. This behavior resembles a proportional–derivative (PD) controller, in which corrective steering depends on the magnitude of the positional error (θ, **Fig. 1A**) and the sign of the derivative term (*dθ/dt*). In a PD controller, corrective output increases when error is growing (*d*θ/*dt* > 0) and decreases when error is shrinking (*d*θ/*dt* < 0); this is useful because it reduces overshooting and improves stability^63^. Notably, however, this anticipatory component only appears when the target is inside the central visual zone. A similar phenomenon has been described in houseflies^15^. Our results help explain why: the central zone pathway (AOTU019) has higher direction selectivity than the peripheral pathway (AOTU025). Moreover, our network model shows how direction-selectivity in the central pathway can improve performance, because it minimizes overshooting. Other insects also show zone-specific direction selectivity, with these zones reflecting each species’ specialized pursuit strategies^15,18,64–68^.

A third factor in adaptive control is forward velocity. Object-directed steering is more vigorous during fast running, and AOTU019 steering drive increases at higher forward speeds. This makes sense: the impact of error in the pursuer’s orientation is increased at higher velocity. Our data also reveal a bidirectional coupling between AOTU019 activity and forward velocity: fast running recruits AOTU019 activity, which drives continued fast running. This positive feedback loop should contribute to the persistence of pursuit behavior^51,69^. By comparison, AOTU025 is less sensitive to forward velocity, and activation of AOTU025 does not further increase forward velocity. This makes good sense: the pursuer should increase their forward velocity only if they have successfully oriented toward the target object, but not if the object is still located in their visual periphery. Among their downstream DNs, DNa03 receives strong input from AOTU019 and far less from AOTU025 (**Fig. 1C, S1A,B**), and because DNa03 activity is associated with slower walking^49,70^, AOTU019 may increase forward velocity by inhibiting DNa03. It is worth noting that many vehicles also use gain scheduling to produce changes in steering control at high forward speeds, although for different reasons^2,63^.

The final factor in adaptive control is behavioral state. Broadly speaking, the effect of social arousal^12^ or attention^14^ on insect visual pursuit is analogous to the effect of attention on primate oculomotor pursuit^4–11^: increased motivation produces more vigorous pursuit performance. Our results suggest that arousal increases pursuit vigor, in part, via the selective recruitment of AOTU019. Depolarization of AOTU019 should inhibit DNa03, which might increase forward velocity^49^. Our findings add to a growing body of evidence that motivated behavioral states can recruit specialized circuits for motor control. For example, larval zebrafish can make more precise eye movements while hunting^71^, due to the recruitment of additional motor neurons^72^. Similarly, flies can execute more precise takeoff maneuvers—if time is available—by selecting a high-precision neural pathway^73^.

Our findings pose several new problems. For example, we do not yet know why AOTU019 is more direction-selective than AOTU025, as both AOTU cell types should logically inherit direction selectivity from LC10a^45^. Also, in a true PD controller, steering should depend on the object’s speed (not just its direction), but AOTU019 responses (like LC10a responses^40^) are not strongly tuned to object speed (**Fig. S4**). Moreover, it is puzzling that behavior is so strongly direction-selective, whereas AOTU019 is only modestly so; it is also puzzling that silencing AOTU019 does not impair behavioral direction-selectivity (**Fig. S7E**). Perhaps behavioral direction selectivity reflects additional contributions from other pathways that detect the direction of optic flow^74–77^.

In the future, it will be interesting to compare circuits for visual pursuit during flight as well as running. AOTU019 makes strong connections onto flight-steering-DNs^56,59,78^(**Fig. S1A**), and it has a remarkably thick cable diameter (**Fig. S5D,E**), which suggests a specialization for the fast processing speeds required for flight. Also, in the future, it will be interesting to investigate the nonvisual inputs to AOTU019 and AOTU025, which seem to influence steering in darkness. The top shared input to AOTU019 and AOTU025 (aside from LC10) comes from a population of inter-hemispheric GABAergic neurons that is positioned to mediate reciprocal inhibition between the right and left AOTU (**Fig. S5C**). These interhemispheric neurons (called AOTU041) are positioned to decorrelate activity in the right and left populations of AOTU cells, which might promote turning with or without a visual stimulus. In addition, AOTU019 is directly downstream from central complex output neurons (i.e., PFL3 cells^70,79^; **Fig. S5C**) which may contribute to steering and forward velocity control, even in darkness.

In summary, our work illustrates two general strategies for implementing adaptive control in a biological feedback system. The first strategy is to create two or more parallel pathways inside a feedback loop, with different gains or control rules in each pathway. Incoming signals can dynamically engage these pathways (alone or together) to produce adaptive control with smooth interpolation. The second strategy is to use internal state variables to recruit specific pathways, or to tune the gain of one pathway. In this study, the key internal state variables are forward velocity commands and arousal states. Together, these strategies allow the system to implement different control rules in different conditions. These principles may offer solutions for the design of mobile engineered systems, which often struggle to achieve the flexibility and adaptability of biological systems^80–82^.

## Resource availability

### Lead contact

Requests for further information and resources should be directed to and will be fulfilled by the lead contact, Rachel Wilson.

### Materials availability

This study did not generate new unique reagents.

## Data and code availability

● All data reported in this paper has been deposited at Zenodo and is publicly available at https://doi.org/10.5281/zenodo.15474937
● All original code for data acquisition, data analysis, and model implementation is available at https://github.com/wilson-lab/CollieWilson.
● Additional information is available from the lead contact upon request.

## Acknowledgements

We thank F. Loesche, M. Reiser, O. Mazor, and P. Gorelik for their assistance with the visual panorama systems; C. Schretter for sharing the aIPg-iLexA driver line; H. Yang for sharing data acquisition and analysis code; A. Bates for sharing connectome and cable width analysis code; F.M. Contini for supporting driver line validation; and the Wilson lab members for their feedback. We thank the Princeton FlyWire team and members of the Allen Institute for Brain Science (supported by BRAIN Initiative grants MH117815 and NS126935 to Murthy and Seung), the Janelia FlyEM team and Cambridge Drosophila Connectomics Group/MRC LMB (supported by HHMI and Wellcome Trust), W.-C.A. Lee and Wilson labs, as well as members of the FlyWire consortium for sharing the connectomes and relevant neuron proofreading and annotation. This research was supported by the Harvard Medical School Research Instrumentation Core (NEI P30 Core Grant for Vision Research EY012196) and the Neurobiology Imaging Facility, and fly stocks were obtained from Bloomington Drosophila Stock Center (NIH P40OD018537). Funding was provided by NSF GRFP grant DGE2140743 (to M.F.C.), NIH F31 grant 1F31NS130782 (to M.F.C.), and NIH U19 grant U19NS104655 (to R.I.W.). M.F.C. was a Stuart H.Q. & Victoria Quan Fellow at Harvard Medical School, and R.I.W. is an HHMI Investigator.

## Author contributions

M.F.C. and R.I.W. designed the study. M.F.C. performed electrophysiology experiments, generated the models, and analyzed the data. C.J., and V.R. conducted behavioral experiments. E.K., C.J., and V.R. generated and described the driver lines used in this study. Q.X.V. proofread neurons in the Flywire-FAFB EM dataset. A.K.M. contributed to pilot experiments, and S.L.H. provided technical support. M.F.C. and R.I.W. wrote the manuscript.

## Supplemental information

**Fig. S1:**
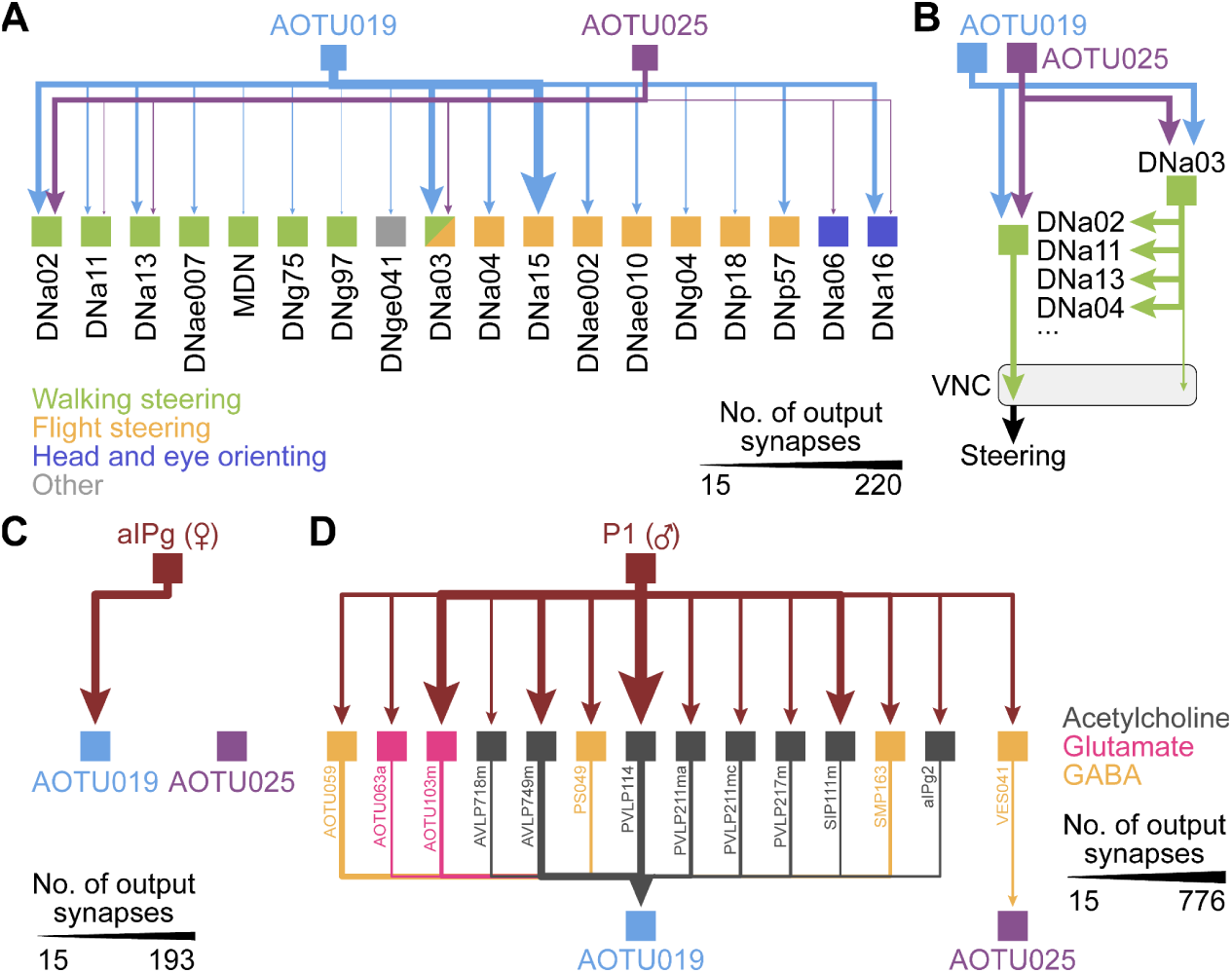
Connectivity of descending neuron outputs and sex-specific arousal inputs. **(A)** Descending neuron (DN) outputs for AOTU019 and AOTU025, with arrow thickness scaled by synapse count (averaged across hemispheres). Several postsynaptic DNs are known to contribute to leg movements during walking (i.e. DNa02^48,57^, DNa11^49^, MDN^50,84,85^, DNa03^49^) or wing movements flight (i.e. DNa03^49^, DNa04^59^, DNa15^86^). Although most DNs have yet to be functionally characterized, recent brain and ventral nerve cord connectivity analyses^24^ suggest that these DNs can be grouped into behaviorally relevant “superclusters,” with most postsynaptic DNs associated with either walking- or flight-related steering. Connectivity (>15 synapses) and superclusters in females from Flywire-BANC. **(B)** DNa02 receives direct input from AOTU019 and AOTU025, and indirect input via DNa03. DNa03 primarily targets other steering-related DNs within the central brain and lacks leg neuromere projections^56^, suggesting an indirect role in steering^49^. **(C)** Shortest paths (>15 synapses) from aIPg cells to AOTU019 and AOTU025 in females. aIPg cells directly target AOTU019 but not AOTU025^29^. pC1d and pC1e cells connect disynaptically to AOTU019 via primarily aIPg cells^29^ and only weakly to AOTU025 through a single inhibitory intermediate (VES041). Synapse counts averaged across hemispheres and pooled across aIPg subtypes (Flywire-FAFB). **(D)** Shortest paths (>15 synapses) from P1 cells to AOTU019 and AOTU025 in males. P1 cells connect disynaptically to AOTU019 via primarily excitatory intermediates and only weakly to AOTU025 through a single inhibitory intermediate. Synapse counts averaged across hemispheres and pooled across P1 subtypes (MaleCNS).

**Fig. S2:**
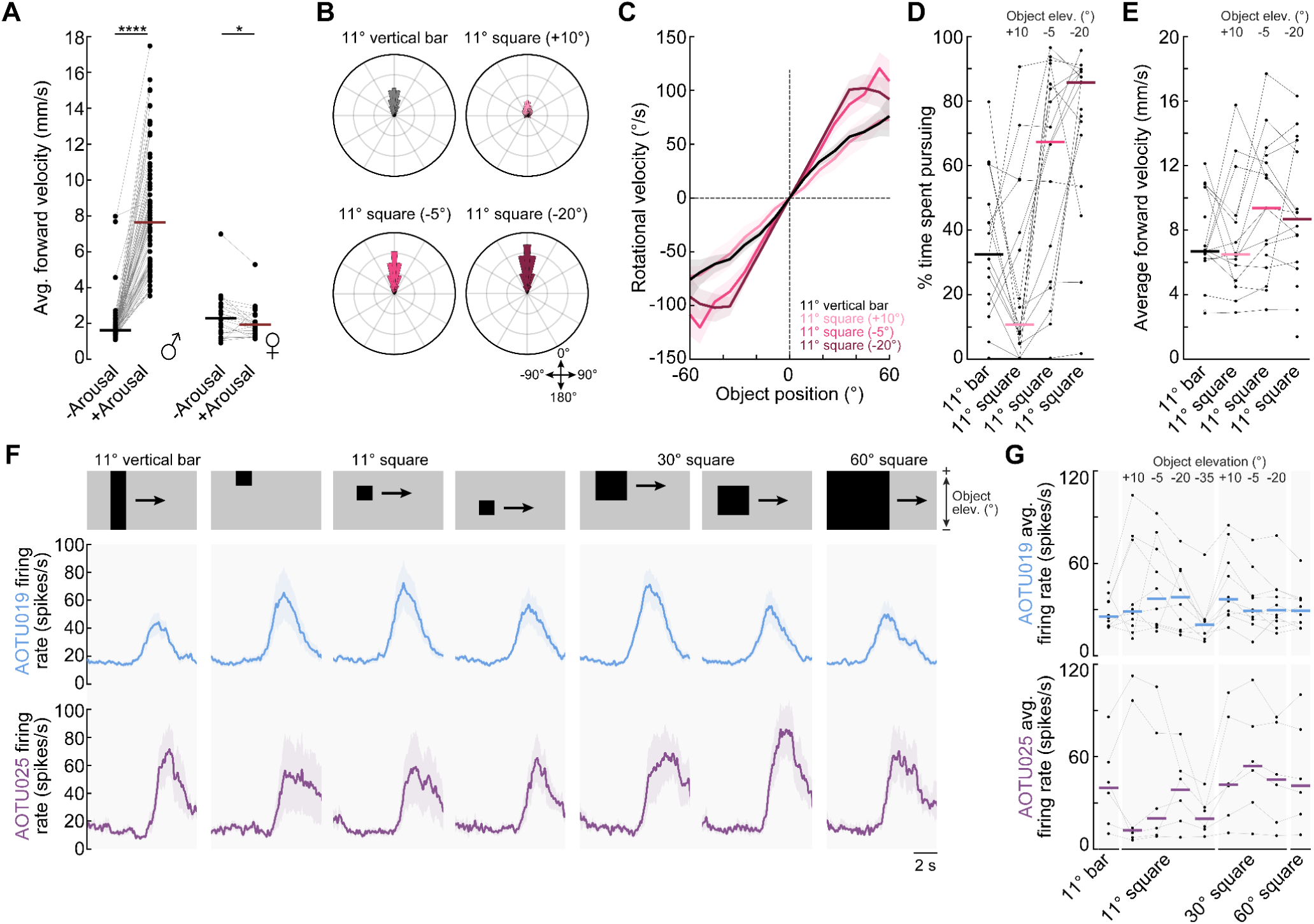
Pursuit behavior and AOTU neuron responses to different visual stimuli. **(A)** Average forward velocity during orientation epochs (object within ±35°) before and during arousal stimulation, shown per fly. Males (P1: n = 75) exhibited a strong increase in forward velocity (p < 0.0001), consistent with “chase” behavior, whereas females (aIPg: n = 21) showed a modest decrease (p < 0.05). Our results would suggest that aIPg only triggers part of the “arousal program”. In females, the cells more functionally analogous to P1 may be pC1d/e cells upstream of aIPg^29,87^. **(B)** Polar histogram of visual object positions for arousal-stimulated males pursuing different visual stimuli in closed loop (n = 17). Data shown as probability densities (range: 0–0.3). Object concentration near the midline did not differ across stimuli (median absolute deviation from 0°, p = 0.332). **(C)** Rotational velocity binned by object position within the frontal field (<60°) during pursuit for different visual stimuli (n = 17). Across the presented stimuli, flies consistently steered toward the visual object. **(D)** Percentage of time males spent pursuing different visual stimuli during arousal stimulation (n = 17). Pursuit engagement differed significantly across stimuli (p < 0.0001). **(E)** Average forward velocity during pursuit epochs for each visual stimulus (n = 17). No significant differences were detected across stimuli (p > 0.05, all post hoc comparisons n.s.). **(F)** Average firing rate responses to visual stimuli during rightward sweeps (–120° to +120°) for AOTU019 (mean ± s.e.m., n = 10) and AOTU025 (n = 7). Stimuli were presented in open loop at 25 °/s (oscillatory triangle waveform) and included a dark vertical bar (11.25°) and dark squares (11.25°, 30°, 60°) at elevations of +10°, –5°, –20°, and –35° relative to the fly’s midline. The vertical bar and 60° square spanned the full vertical extent (60°) of the display. **(G)** Average firing rates during ipsilateral sweeps (0–120°, rightward) for each visual stimulus for AOTU019 (n = 10) and AOTU025 (n = 7). Horizontal lines indicate medians. Responses resembled previously reported LC10 activity^40,45^. Because parts of the dorsal and ventral visual fields were occluded by the treadmill and mounting platform^88–90^, moving bars and squares may have appeared more similar from the fly’s perspective.

**Fig. S3:**
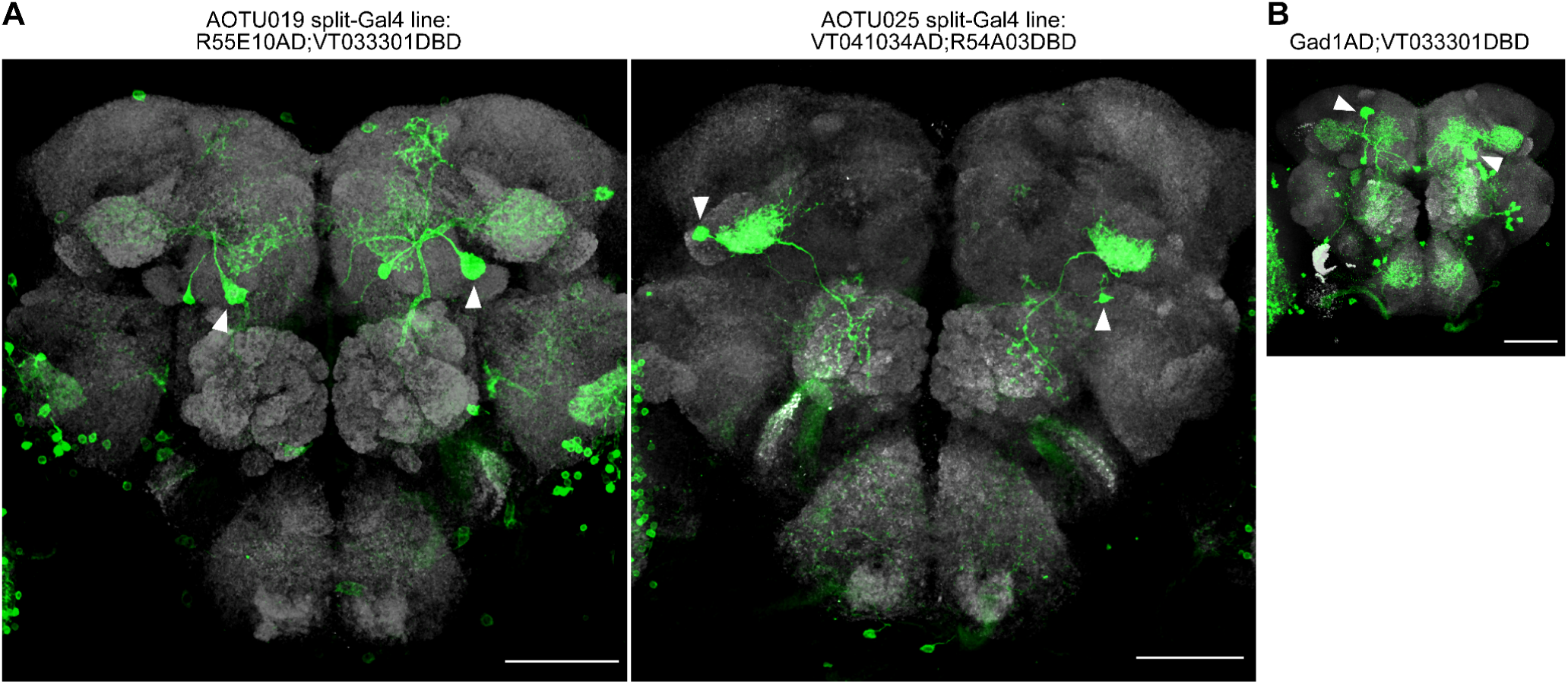
Split-Gal4 lines and neurotransmitter validation. **(A)** Confocal images (maximum z-projections) of split-Gal4 lines targeting AOTU019 and AOTU025, driving GFP expression (green) in the central brain. Neuropil is counterstained with nc82 (gray). Arrowheads indicate left and right cell bodies. The AOTU019 line also labels a small off-target cell near the AOTU019 soma, but its smaller soma size allowed reliable targeting of AOTU019 for recording. Scale bar: 50 µm. **(B)** Validation of AOTU019’s GABAergic identity, as predicted by automated classifications^54^ from Flywire-FAFB and MaleCNS. Confocal image of AOTU019 labeled with the Gad1.AD hemidriver^55^ (Gad1.AD;VT033301.DBD) driving GFP (green). Arrows mark cell bodies; neuropil stained with nc82 (gray). Scale bar: 50 µm.

**Fig. S4:**
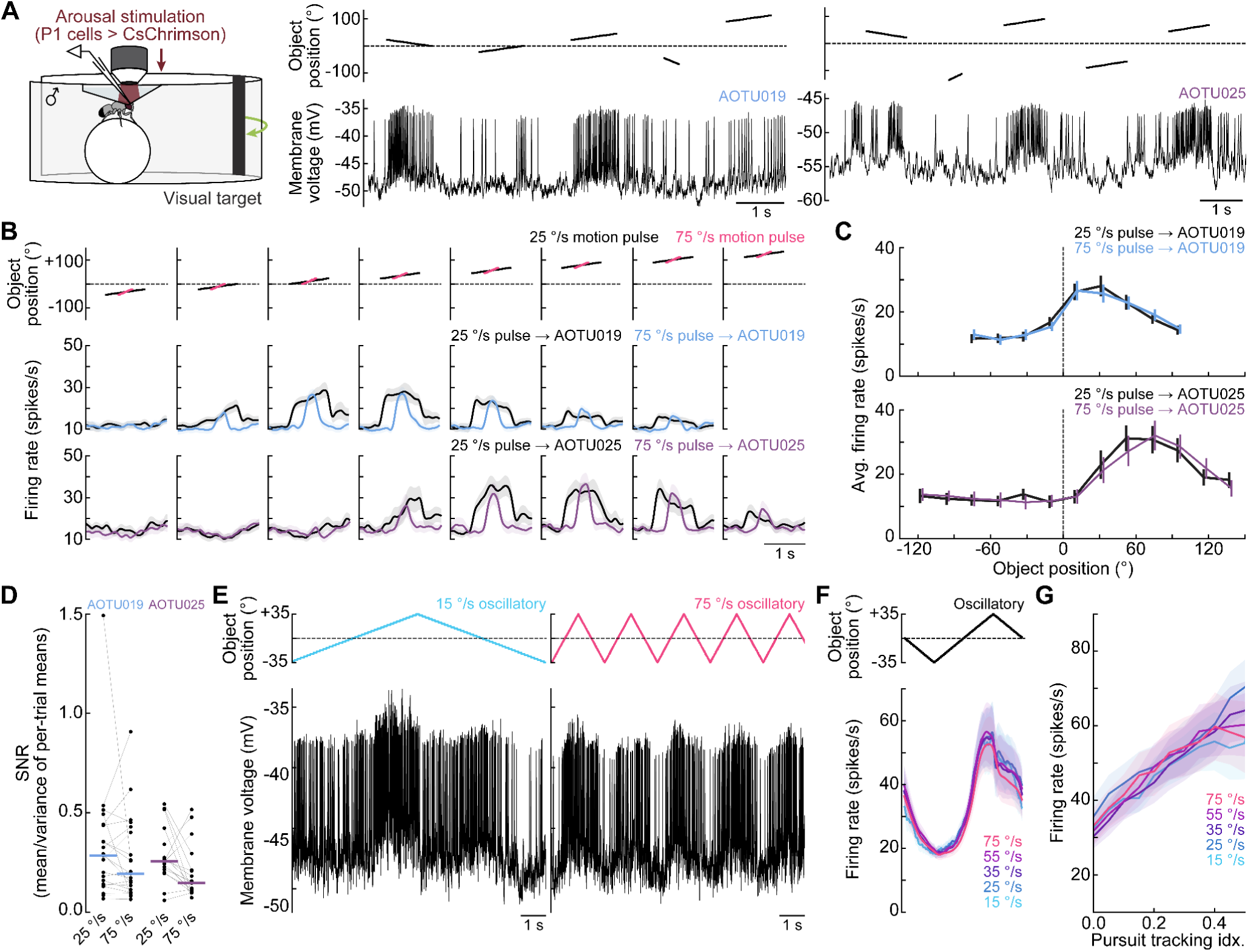
AOTU019 and AOTU025 cells are speed-invariant. **(A)** Schematic of the fly-on-ball setup for arousal-stimulated males. Right: Example recordings from AOTU019 and AOTU025 neurons (right-hemisphere) during P1 stimulation and motion pulse presentations. Visual stimuli spanned 100° of azimuth for AOTU019 and 180° for AOTU025, with each pulse moving 22.5° at 25 or 75 °/s. **(B)** Visual responses to ipsiversive motion pulses at 25 °/s and 75 °/s for AOTU019 (mean ± s.e.m., n = 25) and AOTU025 (n = 22) during stationary epochs in arousal-stimulated males. **(C)** Average firing rates during 25 °/s and 75 °/s motion pulses, plotted at the sweep center for AOTU019 (mean ± s.e.m., n = 25) and AOTU025 (n = 22). No significant differences were observed for either AOTU019 (p = 0.824) or AOTU025 (p = 0.857), consistent with the weak speed tuning of LC10 neurons^40^. **(D)** Signal-to-noise ratio (SNR) for AOTU019 (n = 25) and AOTU025 (n = 22) during stationary epochs in arousal-stimulated males. SNR was calculated as mean firing rate divided by variance across per-trial means, averaged over preferred object positions. No significant differences across speeds (p = 0.440) or cell types (p = 0.176). **(E)** Example AOTU019 recording during oscillatory object motion (±35°, triangle waveform) at 25 °/s (left) and 75 °/s (right) in arousal-stimulated males. Firing increased as the object approached and crossed the midline. **(F)** Average firing rates across oscillatory sweep speeds (15–75 °/s) for AOTU019 (mean ± s.e.m., n = 15) during stationary epochs. Peak firing rates did not differ significantly across speeds (p = 0.991). **(G)** AOTU019 activity binned by pursuit performance^45^ across oscillatory speeds (a.u., mean ± s.e.m., n = 15) in arousal-stimulated males. AOTU019 activity closely tracked pursuit performance, indicating a role in maintaining target fixation.

**Fig. S5:**
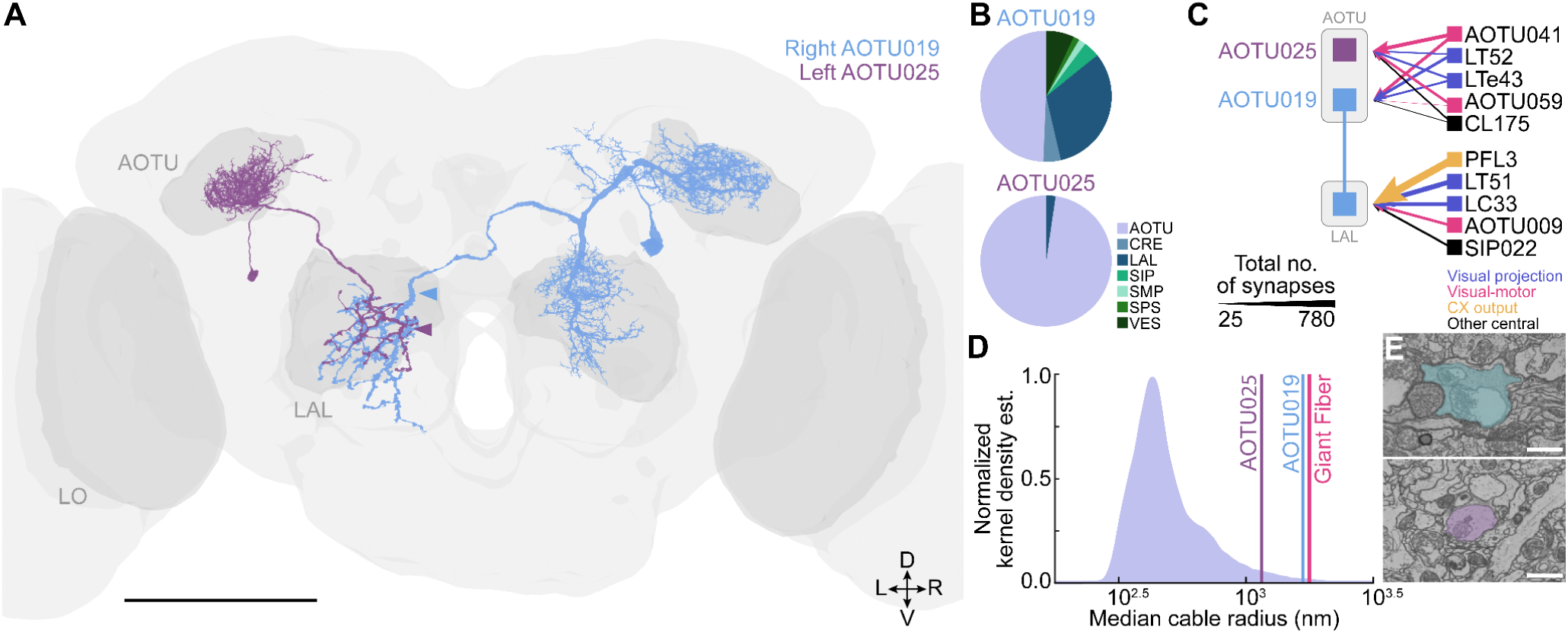
Detailed morphology and non-visual inputs. **(A)** Front-view of the fly brain showing the lobula (LO), anterior optic tubercle (AOTU), and lateral accessory lobe (LAL). Both AOTU019 and AOTU025 have dendrites in the AOTU. AOTU025 projects ipsilaterally (dendrites and axon on the same side), whereas AOTU019 projects contralaterally and also extends dendrites into the LAL and adjacent premotor regions. Premotor input to AOTU019 in the LAL arises from the same hemisphere as its visual input in the AOTU. Neuropil and reconstructions from Flywire-FAFB. Scale bar: 100 µm. **(B)** Distribution of input synapses across neuropil regions (averaged across hemispheres). AOTU025 receives nearly all inputs within the AOTU, whereas AOTU019 also receives substantial input from premotor regions, especially the LAL. The small LAL input to AOTU025 corresponds to axonal synapses. Only regions with ≥50 synapses are shown. Abbreviations: CRE, crepine; SIP, superior intermediate protocerebrum; SMP, superior medial protocerebrum; SPS, superior posterior slope; VES, vest. Connectivity from Flywire-FAFB (females). **(C)** Schematic of top shared inputs (>20 synapses, excluding LC10) to AOTU019 and AOTU025, and top five AOTU019-specific inputs from premotor regions (e.g., LAL). Arrow thickness scales with total synapse count across all cells of a given type (averaged across hemispheres). The central complex (CX) sends steering signals to DNa02 via PFL3 neurons^70,79^, largely via indirect connections through AOTU019^48^. Connectivity from Flywire-FAFB (females). **(D)** Normalized kernel density distribution of median cable (primary dendrite) width across all neurons in Flywire-FAFB. AOTU025 exceeds the 95th percentile, while AOTU019 and the giant fiber exceed the 99th percentile. **(E)** Electron micrograph (EM) cross-sections of AOTU019 (blue, left) and AOTU025 (magenta, right) from Flywire-FAFB. Section planes indicated by arrowheads in **(A)**. Scale bar: 1 µm.

**Fig. S6:**
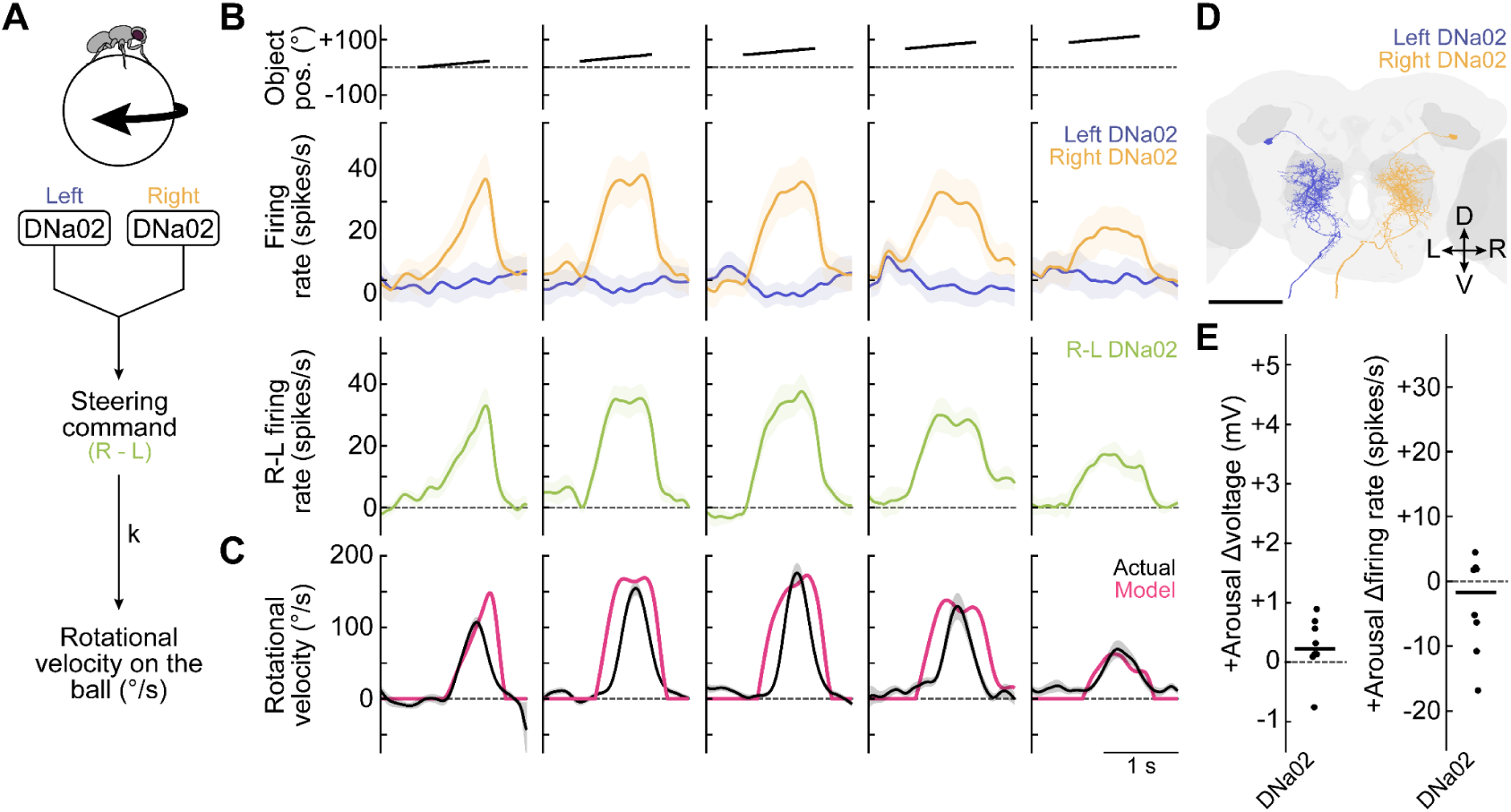
Model of rotational velocity from DNa02 activity. **(A)** Schematic showing how modeled rotational velocity was derived from the right–left difference in DNa02 firing rate. In this model, the R–L activity was integrated over a 100 ms causal window to estimate cumulative drive, then scaled by a gain factor (*k*) and offset by a constant opposing force (*f*) that represents friction. The modeled ball began to move once the cumulative DNa02 activity exceeded this opposing force. **(B)** We estimated the right–left firing rate differences (R–L) for each cell type, assuming mirror-symmetric tuning between hemispheres (see Methods). For each fly, we directly measured the response of the ipsilateral cell (yellow: mean ± s.e.m., n = 11) and estimated the response of its contralateral counterpart by mirroring the visual stimulus across the midline (blue). Under this symmetry assumption, the left cell’s response to a rightward-moving object on the right equals the right cell’s response to a leftward-moving object on the left. We then computed the R-L firing rate difference for every motion pulse stimulus (green), representing the net steering drive contributed by that cell type. This approach follows the “see-saw” steering model (**Fig. 4C**), where steering behavior depends on the difference in activity between the right and left sides. Even small reductions in contralateral responses—such as through inhibition—can substantially increase the overall R–L difference. **(C)** Pink: Modeled rotational velocity derived from right-left firing rate difference of DNa02. Black: Observed rotational velocity (mean ± s.e.m., n = 40). **(D)** Front view of the fly brain showing DNa02 dendrites in premotor regions and ipsilateral projections out of the central brain to the ventral nerve cord, terminating in the leg neuromeres^48,56,57^. Neuropil and cell reconstructions from Flywire-FAFB. Shaded regions: LO, AOTU, and LAL. Scale bar, 100 µm. **(E)** Arousal-induced changes in membrane voltage and firing rate for DNa02 in males (P1: n = 8). Changes in DNa02 activity were not significantly different from zero (p = 0.215; = 0.223). Activity measured during stationary epochs. The 250 ms after activation onset/offset were excluded. Horizontal lines indicate medians.

**Fig. S7:**
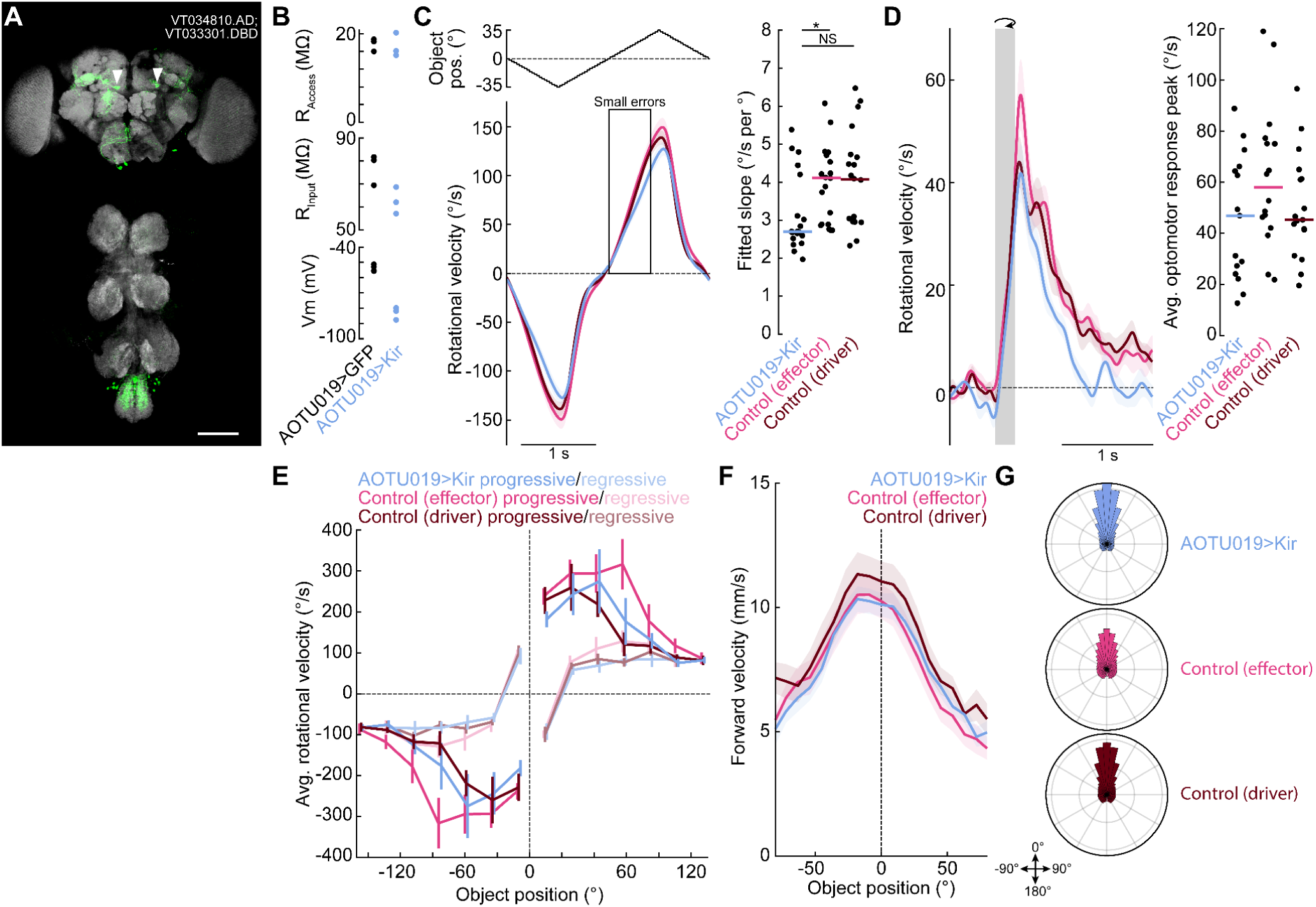
Effects of Kir expression in AOTU019 cells. **(A)** Maximum z-projection of the AOTU019 split-Gal4 line (VT034810.AD; VT033301.DBD) driving GFP (green) in the brain (top) and ventral nerve cord (VNC, bottom). Neuropil labeled with nc82 (gray). This line was used for behavioral experiments due to its sparse expression. Off-target expression was observed in the subesophageal zone (SEZ) and abdominal neuromeres (ANm). Arrows mark AOTU019 cell bodies. Scale bar: 50 µm. **(B)** Validation of Kir2.1-induced hyperpolarization in AOTU019. Whole-cell recordings from Kir2.1 and GFP control flies (n = 3 each) showed ∼30 mV hyperpolarization in Kir2.1-expressing cells. Passive and active properties were measured upon break-in; some Kir2.1 cells required >1 nA current injection to elicit spiking. **(C)** Oscillatory motion (open loop): Turn responses to oscillatory object motion across the front field (mean ± s.e.m., n = 20 each). Right: slope distributions show a significant genotype effect (p < 0.05); AOTU019>Kir differed from the effector control (p < 0.05) and trended toward significance versus the driver control (p = 0.072). Horizontal lines indicate medians. **(D)** Full-field gratings (open loop): Turn responses to clockwise and counterclockwise motion for AOTU019>Kir, effector control, and driver control flies (mean ± s.e.m., n = 20 each). No significant difference in peak responses across genotypes (p = 0.179). **(E)** Motion pulses (open loop): Rotational velocity versus object position (mean ± s.e.m., n = 20 each). As in **Fig. 5A**, flies generally turned toward the object, except near the midline, where turning followed motion direction. Turn responses varied significantly with genotype across positions (genotype-position interaction, p < 0.01), but there was no significant interaction of genotype with direction (p = 0.3748), indicating that AOTU019 is not essential for direction-selective steering near the midline. It is likely that direction-selective steering arises from the combined influence of AOTU019 and other visuo-motor pathways, including optic flow circuits^74–76^. **(F)** Forward velocity binned by object position for AOTU019>Kir (n = 21), effector control (n = 19), and driver control (n = 21) flies (mean ± s.e.m.) during closed-loop pursuit. These curves are similar for all three genotypes, indicating that AOTU019 is not required for the dependence of forward velocity on visual object position. However, it is required for the sharp forward sharp acceleration response when the object sweeps across the midline (**Fig. 7D**). **(G)** Polar histogram of visual object positions during pursuit forAOTU019>Kir (n = 21), effector control (n = 19), and driver control (n = 21) flies. Data shown as probability densities (range: 0–0.11). Genotypes did not differ significantly in the concentration of visual object positions near the midline (median absolute deviation from 0°: p = 0.0993). This shows that other visuo-motor pathways (aside from AOTU019) are sufficient to keep the object near the midline, even though steering vigor is lower when AOTU019 is silenced (**Fig. 7A,B**).

**Fig. S8:**
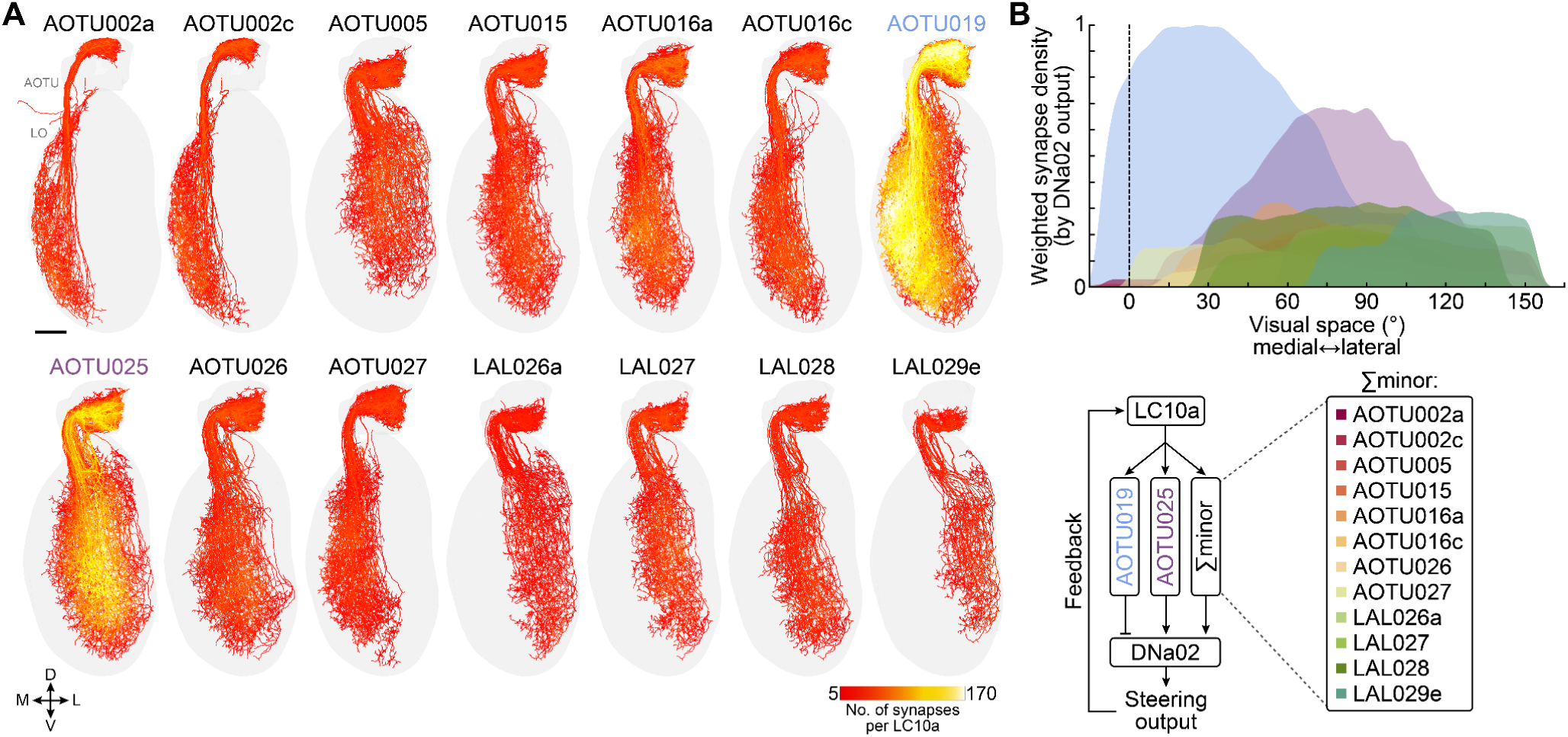
Receptive field estimates and weighted contributions of “minor” AOTU pathways. **(A)** LC10a cells with dendrites in the LO and axons in the AOTU, color-coded by synapse count onto AOTU cell types that both receive direct LC10a input and form direct outputs to DNa02 (a steering-related DN). Neuron skeletons and connectivity from the maleCNS. Scale bar: 20 µm. **(B)** Estimated visual receptive fields for each AOTU cell type, based on LC10a input synapse density along the LO medial–lateral axis in males. Receptive fields were weighted by each cell type’s postsynaptic output strength to DNa02 (**Fig. 1C**). In simulations, values were normalized to a maximum of 1; for instance, the AOTU019→DNa02 connection is ∼5× stronger than the AOTU002a→DNa02 connection, so AOTU002a’s maximum contribution was set to 0.2. Although each “minor” AOTU cell type contributes less individually than AOTU019 or AOTU025, their summed input and output fields likely produce a comparable overall effect. Thus, in the extended dynamical model, AOTU019 and AOTU025 are modeled explicitly, while all other AOTU cell types are combined into a single “minor” unit (see **Fig. 7C**).

**Table S1.** Statistics related to Figures 1-8, S1-8.

| Figure | Test | p-values |
| --- | --- | --- |
| 1E (left) | Paired t-test comparing the median absolute deviation from 0° of object positions before versus during male arousal stimulation | $p = 7.81 \times 10^{-12}$ |
| 1E (right) | Paired t-test comparing the median absolute deviation from 0° of object positions before versus during female arousal stimulation | $p = 7.36 \times 10^{-5}$ |
| 2B (left) | Two-sample t-test for male arousal driven (before vs. during) change in voltage between cell types | $p = 8.99 \times 10^{-7}$ |
| 2B (left) | Two-sample t-test for female arousal driven (before vs. during) change in voltage between cell types | $p = 1.45 \times 10^{-4}$ |
| 2B (right) | Two-sample t-test for male arousal driven (before vs. during) change in firing rate between cell types | $p = 4.42 \times 10^{-6}$ |
| 2B (right) | Two-sample t-test for female arousal driven (before vs. during) change in firing rate between cell types | $p = 3.56 \times 10^{-4}$ |
| 2C | Repeated-measures ANOVA for avg. male visual responses with cell type and arousal as factors | $p(\text{cell type}) = 0.0804$<br>$p(\text{arousal}) = 5.35 \times 10^{-9}$<br>$p(\text{interaction}) = 0.0333$ |
| 2C | Post hoc paired t-tests to assess effect of arousal conditions on avg. male visual responses within each cell type, with Bonferroni correction applied | $p(\text{AOTU019}) = 2.13 \times 10^{-5}$<br>$p(\text{AOTU025}) = 0.0057$ |
| 2C | Repeated-measures ANOVA for avg. female visual responses with cell type and arousal as factors | $p(\text{cell type}) = 0.9889$<br>$p(\text{arousal}) = 0.0001$<br>$p(\text{interaction}) = 0.3086$ |
| 2C | Post hoc paired t-tests to assess effect of arousal conditions on avg. female visual responses within each cell type, with Bonferroni correction applied | $p(\text{AOTU019}) = 0.0046$<br>$p(\text{AOTU025}) = 0.0027$ |
| 3F (top) | Repeated-measures ANOVA for avg. male visual responses with cell type and sweep position as factors | $p(\text{cell type}) = 0.1395$<br>$p(\text{sweep position}) = 0.9591$<br>$p(\text{interaction}) = 0.6711$ |
| 3F (bottom) | Repeated-measures ANOVA for avg. female visual responses with cell type and sweep position as factors | $p(\text{cell type}) = 0.4645$<br>$p(\text{sweep position}) = 0.0138$<br>$p(\text{interaction}) = 0.2123$ |
| 4B (left) | Linear mixed-effects ANOVA for Δrotational or Δforward velocity with AOTU019 current injection (positive, negative, control) as the sole factor | $p(\Delta\text{rotational}) = 3.82 \times 10^{-9}$<br>$p(\Delta\text{forward}) = 6.01 \times 10^{-6}$ |
| 4B (left) | Post hoc Tukey-Kramer comparison for Δrotational velocity was performed to assess AOTU019 current injection (positive, negative, control) differences | $p(\text{control vs. depol.}) = 8.75 \times 10^{-6}$<br>$p(\text{control vs. hyperpol.}) = 0.0344$ |
| 4B (left) | Post hoc Tukey-Kramer comparisons for Δforward velocity was performed to assess AOTU019 current injection (positive, negative, control) differences | $p(\text{control vs. depol.}) = 0.0035$<br>$p(\text{control vs. hyperpol.}) = 0.0661$ |
| 4B (right) | Linear mixed-effects ANOVA for Δrotational or Δforward velocity with AOTU025 current injection (positive, negative, control) as the sole factor | $p(\Delta\text{rotational}) = 1.24 \times 10^{-6}$<br>$p(\Delta\text{forward}) = 0.2458$ |
| 4B (right) | Post hoc Tukey-Kramer comparison for Δrotational velocity to assess AOTU025 current injection (positive, negative, control) differences | $p(\text{control vs. depol.}) = 3.83 \times 10^{-5}$<br>$p(\text{control vs. hyperpol.}) = 0.3621$ |
| 4G (top) | Two-sample t-test for peak rotational correlation between cell types | $p = 0.0088$ |
| 4G (bottom) | Two-sample t-test for peak rotational lag between cell types | $p = 0.0044$ |
| 4H | Repeated-measures ANOVA for binned rotational velocity with cell type and firing rate bin as factors | $p(\text{cell type}) = 0.1557$<br>$p(\text{firing rate bin}) = 1.18 \times 10^{-144}$<br>$p(\text{interaction}) = 0.3876$ |
| 5A | Repeated-measures ANOVA for avg. rotational velocity with sweep position and direction of motion (progressive, regressive) as factors | $p(\text{sweep position}) = 7.35 \times 10^{-14}$<br>$p(\text{direction of motion}) = 7.38 \times 10^{-32}$<br>$p(\text{interaction}) = 5.51 \times 10^{-19}$ |
| 5B (right) | Repeated-measures ANOVA for median DSI with cell type and sweep speed as factors | $p(\text{cell type}) = 0.006$<br>$p(\text{sweep speed}) = 0.456$<br>$p(\text{interaction}) = 0.143$ |
| 5B (right) | Post hoc Tukey-Kramer comparison for median DSI to assess cell type differences at each speed | $p(25^\circ/\text{s}) = 0.0422$<br>$p(75^\circ/\text{s}) = 0.618$ |
| 5C (top) | Repeated-measures ANOVA for AOTU019 avg. visual responses with sweep position and direction of motion (progressive, regressive) as factors | $p(\text{sweep position}) = 6.75 \times 10^{-6}$<br>$p(\text{direction of motion}) = 0.0275$<br>$p(\text{interaction}) = 0.1903$ |
| 5C (bottom) | Repeated-measures ANOVA for AOTU025 avg. visual responses with sweep position and direction of motion (progressive, regressive) as factors | $p(\text{sweep position}) = 1.08 \times 10^{-12}$<br>$p(\text{direction of motion}) = 0.6674$<br>$p(\text{interaction}) = 0.0814$ |
| 6B | Paired t-test comparing the median absolute deviation from $0^\circ$ of object positions during low versus high forward velocity epochs | $p = 2.58 \times 10^{-8}$ |
| 6D | Paired t-test for fitted rotational velocity slopes during low versus high forward velocity epochs | $p = 1.19 \times 10^{-14}$ |
| 6F | Repeated-measures ANOVA with cell type and forward velocity bin as factors | $p(\text{cell type}) = 0.0123$<br>$p(\text{forward velocity bin}) = 2.25 \times 10^{-70}$<br>$p(\text{interaction}) = 1.35 \times 10^{-14}$ |
| 6H (top) | Repeated-measures ANOVA for AOTU019 avg. visual responses with sweep position and forward velocity as factors | $p(\text{sweep position}) = 2.34 \times 10^{-6}$<br>$p(\text{forward velocity}) = 0.00167$<br>$p(\text{interaction}) = 0.821$ |
| 6H (bottom) | Repeated-measures ANOVA for AOTU025 avg. visual responses with sweep position and forward velocity as factors | $p(\text{sweep position}) = 0.331$<br>$p(\text{forward velocity}) = 0.768$<br>$p(\text{interaction}) = 0.753$ |
| 7B | One-way ANOVA for fitted rotational velocity slopes with genotype as the sole factor | $p = 0.008$ |
| 7B | Post hoc Tukey-Kramer comparison for fitted rotational velocity slopes were performed to assess genotype differences | $p(\text{AOTU019} > \text{Kir vs. effector control}) = 0.0092$<br>$p(\text{AOTU019} > \text{Kir vs. driver control}) = 0.0478$<br>$p(\text{effector control vs. driver control}) = 0.7657$ |
| 7D | Repeated-measures ANOVA for forward velocity response with genotype and object position as factors | $p(\text{genotype}) = 0.4331$<br>$p(\text{object position}) = 1.30 \times 10^{-61}$<br>$p(\text{interaction}) = 1.44 \times 10^{-12}$ |
| 7D | Post hoc pairwise comparison of forward velocity response tuning curves between genotypes using mixed-effects model | $p(\text{AOTU019} > \text{Kir vs. effector control}) = 2.60 \times 10^{-14}$<br>$p(\text{AOTU019} > \text{Kir vs. driver control}) = 8.46 \times 10^{-14}$<br>$p(\text{effector control vs. driver control}) = 0.9994$ |
| 7F | One-way ANOVA for avg. forward velocity during pursuit with genotype as the sole factor | $p = 0.4765$ |
| 7G | Repeated-measures ANOVA for percentage of time spent pursuing with genotype and arousal as factors | $p(\text{genotype}) = 0.99998$<br>$p(\text{arousal}) = 4.26 \times 10^{-18}$<br>$p(\text{interaction}) = 0.0807$ |
| 7G | Post hoc paired t-tests to assess effect of arousal conditions on time spent pursuing within each genotype, with Bonferroni correction applied | $p(\text{AOTU019} > \text{Kir}) = 1.91 \times 10^{-7}$<br>$p(\text{effector control}) = 7.14 \times 10^{-7}$<br>$p(\text{driver control}) = 6.31 \times 10^{-8}$ |
| 8B (top) | Repeated-measures ANOVA for the effect of arousal and visual condition on AOTU019 peak correlation with rotational velocity | $p(\text{arousal}) = 4.47 \times 10^{-5}$<br>$p(\text{visual condition}) = 0.1410$ |
| 8B (top) | Post-hoc pairwise comparison of AOTU019 peak correlation with rotational velocity between arousal conditions in darkness using mixed-effects model. | $p = 4.97 \times 10^{-5}$ |
| 8B (bottom) | Repeated-measures ANOVA for the effect of arousal and visual condition on AOTU019 peak rotational lag | $p(\text{arousal}) = 0.0084$<br>$p(\text{visual condition}) = 0.1934$ |
| 8B (bottom) | Post-hoc pairwise comparison of AOTU019 peak rotational lag between arousal conditions in darkness using mixed-effects model. | $p = 0.0053$ |
| 8B (top) | Repeated-measures ANOVA for the effect of arousal and visual condition on AOTU025 peak correlation with rotational velocity | $p(\text{arousal}) = 0.0021$<br>$p(\text{visual condition}) = 0.8604$ |
| 8B (top) | Post-hoc pairwise comparison of AOTU025 peak correlation with rotational velocity between arousal conditions in darkness using mixed-effects model. | $p = 9.52 \times 10^{-4}$ |
| 8B (bottom) | Repeated-measures ANOVA for the effect of arousal and visual condition on AOTU025 peak rotational lag | $p(\text{arousal}) = 0.2451$<br>$p(\text{visual condition}) = 0.2847$ |
| S2A (left) | Paired t-test for avg. forward velocity during orientation epochs before and during male arousal stimulation | $p = 2.57 \times 10^{-27}$ |
| S2A (right) | Paired t-test for avg. forward velocity during orientation epochs before and during female arousal stimulation | $p = 0.0390$ |
| S2B | Repeated-measures ANOVA for median absolute deviation from $0^\circ$ of object positions with visual stimulus presented as the sole factor | $p = 0.3317$ |
| S2D | Repeated-measures ANOVA for percentage of time spent pursuing with visual stimulus presented as the sole factor | $p = 1.16 \times 10^{-7}$ |
| S2D | Post hoc Tukey-Kramer comparison for percentage of time spent pursuing to assess visual stimulus differences | $p(\text{bar vs. square } +10^\circ) = 0.6815$<br>$p(\text{bar vs. square } -5^\circ) = 0.0258$<br>$p(\text{bar vs. square } -20^\circ) = 7.26 \times 10^{-5}$<br>$p(\text{square } +10^\circ \text{ vs. square } -5^\circ) = 3.18 \times 10^{-4}$<br>$p(\text{square } +10^\circ \text{ vs. square } -20^\circ) = 3.76 \times 10^{-7}$<br>$p(\text{square } -5^\circ \text{ vs. square } -20^\circ) = 0.3878$ |
| S2E | Repeated-measures ANOVA for avg. forward walking speed with visual stimulus presented as the sole factor | $p = 0.0488$ |
| S2E | Post hoc Tukey-Kramer comparison for forward walking speed to assess visual stimulus differences | $p(\text{bar vs. square } +10^\circ) = 0.9999$<br>$p(\text{bar vs. square } -5^\circ) = 0.2032$<br>$p(\text{bar vs. square } -20^\circ) = 0.2101$<br>$p(\text{square } +10^\circ \text{ vs. square } -5^\circ) = 0.2836$<br>$p(\text{square } +10^\circ \text{ vs. square } -20^\circ) = 0.2923$<br>$p(\text{square } -5^\circ \text{ vs. square } -20^\circ) = 1$ |
| S4C (top) | Repeated-measures ANOVA for AOTU019 with sweep position and sweep speed as factors | $p(\text{position}) = 1.829 \times 10^{-21}$<br>$p(\text{speed}) = 0.8956$<br>$p(\text{interaction}) = 0.9942$ |
| S4C (bottom) | Repeated-measures ANOVA for AOTU025 with sweep position and sweep speed as factors | $p(\text{position}) = 2.01 \times 10^{-15}$<br>$p(\text{speed}) = 0.9200$<br>$p(\text{interaction}) = 0.99998$ |
| S4D | Repeated-measures ANOVA for SNR with cell type and object speed as factors | $p(\text{cell type}) = 0.1756$<br>$p(\text{speed}) = 0.4401$<br>$p(\text{interaction}) = 0.4013$ |
| S4F | Repeated-measures ANOVA for peak firing rate with object speed as a fixed effect | p = 0.9909 |
| S6E | One-sample t-test for for male arousal driven (before vs. during) change in voltage for DNa02 | p = 0.2145 |
| S6E | One-sample t-test for for male arousal driven (before vs. during) change in firing rate for DNa02 | p = 0.2232 |
| S7C (right) | One-way ANOVA for fitted rotational velocity slopes with genotype as the sole factor | p = 0.0232 |
| S7C (right) | Post hoc Tukey-Kramer comparisons were performed to assess genotype differences. | p(AOTU019>Kir vs. effector control) = 0.0294<br>p(AOTU019>Kir vs. driver control) = 0.0724<br>p(effector control vs. driver control) = 0.9227 |
| S7D | One-way ANOVA for average optomotor response peak with genotype as the sole factor | p = 0.1786 |
| S7E | Repeated-measures ANOVA for avg. rotational velocity with genotype, object position, and sweep direction as factors | p(position) = $1.07 \times 10^{-6}$<br>p(direction) = $3.12 \times 10^{-12}$<br>p(genotype) = 0.4084<br>p(position x direction) = $3.47 \times 10^{-7}$<br>p(position x genotype) = 0.0019<br>p(direction x genotype) = 0.3748<br>p(position x direction x genotype) = 0.4174 |
| S7E | One-way ANOVA comparing the median absolute deviation from 0° of object positions with genotype as the sole factor | p = 0.0993 |
| Methods | Repeated-measures ANOVA for fixation time with genotype and steering ratio as factors | p(genotype) = 0.0224<br>p(ratio) = 0.3588 |
| Methods | Repeated-measures ANOVA for peak visual-motor lag with genotype and steering ratio as factors | p(genotype) = 0.6814<br>p(ratio) = 0.7396 |
| Methods | Repeated-measures ANOVA for fitted rotational velocity slopes with genotype and ratio as factors | p(genotype) = 0.0009<br>p(ratio) = 0.8594 |

## Highlights

● Parallel pathways detect a visual object in the central versus peripheral visual field
● The pathway for objects in the central visual field has adjustable gain
● In this pathway, gain depends on the direction of object motion and running speed
● This pathway is recruited by arousal and it increases the vigor of visual pursuit

## eTOC blurb

Collie et al. show how adaptive control in *Drosophila* visual pursuit arises from two parallel pathways: one steers objects from the periphery toward the center, while the other maintains central fixation and modulates gain based on object motion and running speed. This architecture supports vigorous pursuit while maintaining feedback stability.

## STAR Methods

### EXPERIMENTAL MODEL AND STUDY PARTICIPANT DETAILS

Flies expressing CsChrimson^84^ were reared on cornmeal-molasses food (Archon Scientific) mixed with 100 µl all-*trans*-retinal (Sigma; 17 mM in ethanol) and wrapped in light-blocking foil to prevent photo-conversion. Flies not expressing CsChrimson were reared on standard cornmeal-molasses food. All flies were maintained in an incubator (Darwin Chambers) on a 12 h light:12 h dark cycle at 25 °C, 50-70% relative humidity. All experimental flies had at least one wild-type endogenous copy of the white (*w*) gene, represented by *w*+ in the genotypes below. Genotypes were as follows:

**Fig. 1, 4-6, S2, S6:**

Behavioral measurements in intact flies with P1 activation:

*w+; P{VT034810-p65.AD}attP40/P{GMR71G01-lexA}attP40;*

*P{VT033301-GAL4.DBD}attP2/PBac{13XLexAop2-IVS-CsChrimson.tdTomato}VK00005*

*w+; +/P{GMR71G01-lexA}attP40; +/P{pJFRC49-10xUAS-IVS-eGFP::Kir2.1}attP2*

*PBac{13XLexAop2-IVS-CsChrimson.tdTomato}VK00005*

**Fig. 1, S2:**

Behavioral measurements in intact flies with aIPg activation:

*w+ P{10XUAS-IVS-mCD8::GFP}su(Hw)attP8/w+; P{R55E10-p65.AD}attP40/P{R72C11-ilexA}JK22C;*

*P{VT033301-GAL4.DBD}attP2/PBac{13XLexAop2-IVS-CsChrimson.tdTomato}VK00005*

*w+ P{10XUAS-IVS-mCD8::GFP}su(Hw)attP8/w+; P{VT041034-p65.AD}attP40/P{R72C11-ilexA}JK22C;*

*P{R54A03-GAL4.DBD}attP2/PBac{13XLexAop2-IVS-CsChrimson.tdTomato}VK00005*

**Fig. 2-6, 8, S2-4:**

AOTU019 recording with P1 activation:

*w+ P{10XUAS-IVS-mCD8::GFP}su(Hw)attP8; P{R55E10-p65.AD}attP40/P{GMR71G01-lexA}attP40;*

*P{VT033301-GAL4.DBD}attP2/PBac{13XLexAop2-IVS-CsChrimson.tdTomato}VK00005*

AOTU025 recording with P1 activation:

*w+ P{10XUAS-IVS-mCD8::GFP}su(Hw)attP8; P{VT041034-p65.AD}attP40/P{GMR71G01-lexA}attP40;*

*P{R54A03-GAL4.DBD}attP2/PBac{13XLexAop2-IVS-CsChrimson.tdTomato}VK00005*

**Fig. 2, 3:**

AOTU019 recording with aIPg activation:

*w+ P{10XUAS-IVS-mCD8::GFP}su(Hw)attP8/w+; P{R55E10-p65.AD}attP40/P{R72C11-ilexA}JK22C;*

*P{VT033301-GAL4.DBD}attP2/PBac{13XLexAop2-IVS-CsChrimson.tdTomato}VK00005*

AOTU025 recording with aIPg activation:

*w+ P{10XUAS-IVS-mCD8::GFP}su(Hw)attP8/w+; P{VT041034-p65.AD}attP40/P{R72C11-ilexA}JK22C;*

*P{R54A03-GAL4.DBD}attP2/PBac{13XLexAop2-IVS-CsChrimson.tdTomato}VK00005*

**Fig. 4, S6:**

DNa02 recording with P1 cell activation:

*w+; P{20XUAS-IVS-mCD8::GFP}attP40/P{GMR71G01-lexA}attP40;*

*P{GMR87D07-GAL4}attP2/PBac{13XLexAop2-IVS-CsChrimson.tdTomato}VK00005*

**Fig. 7, S7:**

AOTU019 cells expressing Kir2.1 with P1 activation:

*w+; P{VT034810-p65.AD}attP40/P{GMR71G01-lexA}attP40;*

*P{VT033301-GAL4.DBD}attP2/P{pJFRC49-10xUAS-IVS-eGFP::Kir2.1}attP2*

*PBac{13XLexAop2-IVS-CsChrimson.tdTomato}VK00005*

AOTU019 driver control with P1 activation:

*w+; P{VT034810-p65.AD}attP40/P{GMR71G01-lexA}attP40;*

*P{VT033301-GAL4.DBD}attP2/PBac{13XLexAop2-IVS-CsChrimson.tdTomato}VK00005*

Effector control with P1 activation:

*w+; +/P{GMR71G01-lexA}attP40; +/P{pJFRC49-10xUAS-IVS-eGFP::Kir2.1}attP2*

*PBac{13XLexAop2-IVS-CsChrimson.tdTomato}VK00005*

**Fig. S3:**

AOTU019 and Gad1 line:

*w+; P{20XUAS-IVS-mCD8::GFP}attP40/+;*

*Mi{Trojan-p65AD.2}Gad1[MI09277-Tp65AD.2]/P{VT033301-GAL4.DBD}attP2*

**Fig. S7:**

AOTU019 cells expressing GFP and Kir2.1:

*w+; P{VT034810-p65.AD}attP40/P{20XUAS-IVS-mCD8::GFP}attP40;*

*P{VT033301-GAL4.DBD}attP2/P{pJFRC49-10xUAS-IVS-eGFP::Kir2.1}attP2*

AOTU019 cells expressing GFP:

*w+; P{VT034810-p65.AD}attP40/P{20XUAS-IVS-mCD8::GFP}attP40; P{VT033301-GAL4.DBD}attP2/+*

### Origins of transgenic stocks

The following stocks were obtained from the Bloomington Drosophila Stock Center (BDSC): split-Gal4 hemidrivers to target AOTU019 (BDSC_69919, BDSC_71336, BDSC_73717), split-GAL4 hemidrivers to target AOTU025 (BDSC_73987, BDSC_75635, BDSC_68804), *P{GMR87D07-GAL4}attP2* to target DNa02^56^ (BDSC_48391), *P{GMR71G01-lexA}attP40* to target P1^44^ (BDSC_54733), split-Gal4 AD hemidriver expressed in Gad1+ neurons^55^ (BDSC_60322), *PBac{13XLexAop2-IVS-CsChrimson.tdTomato}VK00005* (BDSC_82183), and *P{20XUAS-IVS-mCD8::GFP}attP2* (BDSC_32194). *P{pJFRC49-10xUAS-IVS-eGFP::Kir2.1}attP2* flies were a gift from G. Card^73^, *P{R72C11-ilexA}JK22C* flies to target aIPg were a gift from C. Schretter^53^, and *w*+ recombined *P{10XUAS-IVS-mCD8::GFP}su(Hw)attP8* and isogenized *P{20XUAS-IVS-mCD8::GFP}attP40* flies were gifts from H. Yang^57^. All wild-type (+) chromosomes were from the isoD1 background (derived from OregonR); isoD1 flies were a gift from T. Clandinin^85^. The recombined *P{pJFRC49-10xUAS-IVS-eGFP::Kir2.1}attP2 PBac{13XLexAop2-IVS-CsChrimson.tdTomato}VK00005* was generated in-house. The isogenized AOTU019 split-Gal4 stocks (backcrossed for five generations into the isoD1 background) were also generated in-house.

We validated the expression of the constructed AOTU019 and AOTU025 split-Gal4 lines using immunohistochemical anti-GFP staining and also by comparison against available Multi-Color-Flip-Out (MCFO) image libraries^86,87^.

## METHOD DETAILS

### Fly preparation and dissection

Flies were collected on CO₂ and had their wings clipped (to reduce grooming and encourage walking on the ball) the day before experiments, and were housed overnight at low density (≤8 flies per vial) in vials of cornmeal molasses food supplemented with all-*trans*-retinal. All experiments used flies 16–30 h post-eclosion without circadian timing restrictions. To prepare an experiment, flies were briefly cold-anesthetized and fit into an inverted pyramid-shaped platform (black Delrin, Protolabs). This platform design minimized eye occlusion and maximized body and leg mobility. The fly’s head and thorax were secured with UV-curable glue (Loctite AA 3972, Henkel), cured with UV light pulses (LED-200, Electro-Lite Co.). For behavioral experiments on intact flies, flies were secured into the platform, but no further dissection was performed. For electrophysiology experiments, after the fly was secured in the platform, the proboscis was immobilized by severing the lateral muscles and securing the appendage with UV-curable glue while the fly was cold-anesthetized (to reduce brain movement). After several minutes of recovery time, a small window was cut in the dorsal cuticle under oxygenated saline. Tracheoles, fat, and perineurial glia sheath were removed. Muscle 16 (and often the esophagus) was severed to further reduce brain movement. Prior to data collection, all flies were given a 20-minute acclimation period on the spherical treadmill with the visual cue rotating in closed loop with the fly’s rotation.

### Experimental recording setups

Electrophysiology experiments used a semi-custom upright microscope with a motorized base (Thorlabs), epifluorescence turret (BX51, Olympus), 40× water immersion objective (LUMPlanFLN 40×W, Olympus), and CCD Monochrome Camera (Retiga ELECTRO, Teledyne Photometrics). The substage optics were removed to fit the treadmill and visual panorama. The brain was back-illuminated with a far-red LED (M780L3, Thorlabs). For GFP or CsChrimson excitation, a 100 W Hg arc lamp (U-LH100HG, Olympus), controlled by an optical shutter, provided blue (480 nm) or red (620-660 nm) light through the objective, respectively. Red light was dimmed using neutral density filters. The microscope was enclosed in a Faraday cage and blackout material, and mounted on an air-supported table for vibration isolation.

Patch-clamp recordings were made using an Axopatch 200B amplifier (Molecular Devices), low-pass filtered at 5 kHz, and acquired at 20 kHz. Oxygenated saline (28-30 °C) superfused the brain during experiments. Saline contained (in mM): 103 NaCl, 3 KCl, 5 TES, 8 trehalose, 10 glucose, 26 NaHCO₃, 1 NaH₂PO₄, 1.5 CaCl₂, and 4 MgCl₂ (osmolarity adjusted to 270–273 mOsm, pH ∼7.3). Remaining perineural sheath or cell bodies were cleared using saline-filled “cleaning pipettes” pulled from thin-wall capillaries (TW150F-3, World Precision Instruments). Cell recordings were made using patch pipettes (6-12 MΩ) pulled from filamented borosilicate glass (B150-86-7.5HP, Sutter Instrument) and filled with 0.22-µm filtered internal solution, containing (in mM): 140 potassium aspartate, 10 HEPES, 4 MgATP, 0.5 Na₃GTP, 1 EGTA, 1 KCl, and 13 biocytin hydrazide (pH 7.3, osmolarity adjusted to 268 mOsm). All data was acquired at 20 kHz (NiDAQ PCIe-6251, National Instruments).

### Spherical treadmill and locomotion measurement

An air-cushioned spherical treadmill equipped with a machine vision system was used to track the animal’s intended movements. The treadmill consisted of a 9 mm diameter foam ball (FR-4615, General Plastics) floated in a custom concave holder (ABS Black, Protolabs) using medical-grade breathing air (Med-Tech) regulated via an inline flow meter (Cole Parmer). For tracking, the ball was marked with a high-contrast pattern and illuminated by a far-red LED (M780L3, Thorlabs). A CMOS camera (GS3-U3-51S5M, Teledyne FLIR) with an adjustable zoom lens (M7528-MP, Computar) captured ball movement at 60 Hz. FicTrac v2.1^88^ tracked ball position, with a custom Python script for transmitting forward, sideways, rotational, and gain-modified rotational displacements to an analogue output device (PhidgetAnalog 4-Output), acquired at 20 kHz (NiDAQ PCIe-6251, National Instruments). In “virtual reality” experiments (i.e., where specifically reported), the visual cue rotated in closed loop with the fly’s rotation on the spherical treadmill.

### Visual panorama

Visual stimuli were displayed using a circular panorama of modular 16×16 pixel LED panels^89^, with each pixel subtending ∼1.875° of the animal’s visual field and covering 60° in height and 330° in azimuth. The rear column (30°) was removed for ball illumination and tracking equipment. Green LEDs (525 nm) were used for electrophysiology and blue LEDs (465 nm) for experiments focused exclusively on behavioral measurements. Diffuser film (SXF-0600, Decorative Films) and gel filters (RE131 and RE071, Rosco) reduced reflections and intensity. For electrophysiology experiments, the panorama was grounded with fine copper mesh. The panorama, controlled at 500 Hz using a reconfigurable I/O device (PCIe-7842R, National Instruments) and display tools from M. Reiser (https://reiserlab.github.io/Modular-LED-Display/G4/), provided real-time output of visual cue position and gain-modified displacement via a breakout box (CB-68LPR, National Instruments) to the data acquisition card.

For all main experiments with visual object pursuit, the visual stimulus was a dark vertical bar (11.25°×60°) on a bright background. For additional experiments testing neuronal and behavioral responses to different visual objects, stimuli included a dark vertical bar (11.25°×60°) and dark squares of varying sizes (11.25°×11.25°, 30°×30°, and 60°×60°) on a bright background. Squares were presented at four elevations, defined by the position of their top edge (approximately [°]: +10, –5, –20, and –35), relative to 0° directly ahead of the fly. For optomotor response experiments, a full-field square wave grating with a 15° period was used. In experiments without a visual object (i.e., darkness), the panorama was uniformly illuminated at low intensity.

### Visual stimuli (open-loop and closed-loop)

During experiments where the visual object moved in a fixed pattern, irrespective of the fly’s own movements (“open loop”), we presented: (1) an object appearing randomly every 3 s, sweeping 22.5° at 25 or 75°/s, then disappearing (“motion pulses”), with motion paths tiling the front 100 or 180° of the visual field, (2) an object moving back and forth across the visual field following a triangle wave pattern (“oscillating object”) with an amplitude of ±35° at speeds of 15, 25, 35, 55, or 75°/s (for object speed experiments) or ±120° at 25°/s (for object shape/size experiments), and (3) full-field square-wave vertical gratings alternating between clockwise and counterclockwise at 75°/s (“optomotor stimuli”).

During experiments where the visual object rotated around the panorama in “closed loop”, the object’s azimuthal position was controlled by the spherical treadmill’s rotational displacement: when the spherical treadmill rotated to the right (signaling an attempted left turn by the fly), the object also rotated right, and vice versa. Every 30 s, the visual object “jumped” 60° left or right relative to the object’s current position, and at each time step, we also added random motion to the visual object; these perturbations challenged the fly pursuit system to keep the object near the fly’s visual midline. The random motion was drawn from a standard normal distribution (mean = 0, SD = 1), low-pass filtered at 2 Hz (−3 dB corner frequency, Butterworth filter), then mean-subtracted and scaled to have a standard deviation of 0.57° (0.010 radians).

In closed loop experiments silencing AOTU019, the ratio of object rotation to spherical treadmill rotation was varied from 0.95× to 2×, with a ratio randomized at the start of each trial. Varying the object ratio was intended to challenge the male’s ability to correct his trajectory during pursuit, however, we found no significant effect of this ratio on the percentage of time the male spent pursuing the object (p = 0.3588), the estimated lag between the visual object’s position and the male’s rotational velocity (p = 0.7396), or the fitted slope of the function relating rotational velocity to object position (p = 0.8594). Thus, in **Figures 7** and **S7F,G** we pooled data across ratios per fly. In **Figure 1E** and **S2A** comparing behavior before and after arousal stimulation, only 0.95× ratio trials were used to match the non-arousal condition. In all other pursuit experiments (i.e., male fly pursuit of different visual stimuli in **Fig. S2B-E** or female fly pursuit of a moving visual target in **Fig. 1E, S2A**), the ratio of object rotation was fixed at 0.95×.

### Behavioral observations in intact flies

For experiments where we tested the effect of Kir2.1 expression, the experimenter was blinded to genotype during data collection and genotypes were randomly interleaved. Flies were secured into our standard fly-restraining platform (described above), but no further dissection was performed. The fly and the visual panorama were placed inside an enclosure warmed to 25-30°C with a fan heater. The enclosure was covered in blackout material to reduce visual disruptions and mounted on sorbothane feet (AV4, Thorlabs) to reduce vibrations. For CsChrimson excitation, a red LED (M660L4, Thorlabs) was positioned above the fly and controlled by a LED driver (LEDD1B, Thorlabs). For CsChrimson-expressing P1 (40-55 µW/mm² at 600 nm) or aIPg cells (15 µW/mm² at 600 nm), we delivered optostimulation for the full trial duration. All data was acquired at 20 kHz (NiDAQ PCIe-6251, National Instruments).

### Experiment trial structure

To drive “arousal” (i.e., motivate visual object pursuit), we optogenetically stimulated CsChrimson-expressing P1 (4 µW/mm² at 600 nm) or aIPg (1.6 µW/mm² at 600 nm) cells with red light delivered for the full trial duration. Although both P1 and aIPg activation increase visuomotor engagement, P1 cells in males are not directly analogous to aIPg cells. P1 cells are likely more comparable to upstream pC1d/e cells in females^29,90^.

For electrophysiology, experiments routinely began with alternating light pulses (15 s off, 15 s on) to stimulate P1/aIPg cells and measure baseline changes in membrane potential and firing rate without visual stimuli. For motion pulse experiments, pulses at different positions and velocities were presented in a pseudo-randomized order. For oscillating object experiments, different object speeds were also presented in pseudo-randomized order. Trials without arousal stimulation, using a 35°/s object, were randomly interleaved between oscillating object trials. Between motion pulse and oscillating object trials, flies were given 5-10 s breaks without optostimulation.

To manipulate cell activity, brief current injections (1 s in duration every 5 s) were delivered via the recording electrode to depolarize (+100 pA) or hyperpolarize (−75 pA) AOTU019, and depolarize (+20-40 pA) or hyperpolarize (−15-20 pA) AOTU025. Current magnitudes were selected to elicit comparable changes in firing rate across cell types. Generally, hyperpolarization largely or completely suppressed spiking, while depolarization increased firing rates by 30–50 spikes/s. Here, the fly walked in darkness and optostimulation was delivered for the full trial duration. Step order was randomized at the start of each trial.

### Immunohistochemistry

Brains were dissected 1–2 days post-eclosion in external saline, fixed in 4% paraformaldehyde (Electron Microscopy Sciences) in PBS (Thermo Fisher Scientific) for 15 min at room temperature, and washed with PBS. After a 20-min block in 5% normal goat serum (Sigma-Aldrich) in PBS with 0.44% Triton-X (Sigma-Aldrich), samples were incubated ∼24 h with primary antibodies in blocking solution, washed with PBS, and incubated ∼24 h with secondary antibodies. Brains were rinsed, mounted in antifade medium (Vectashield, Vector Laboratories), and imaged on a Leica confocal microscope (SPE, SP8X, or Stellaris) with a 20× or 40× oil-immersion objective. Image stacks included 50–200 z-slices (1 µm depth, 1,024×1,024 pixels, 3 line averages). For Gal4 and lexA expression, primary antibodies were chicken anti-GFP (1:1,000, Abcam), rabbit anti-RFP (1:100, Abcam), and mouse anti-Bruchpilot (1:30, nc82, DSHB). Secondary antibodies included Alexa Fluor 488 anti-chicken, Alexa Fluor 568 anti-rabbit, and Alexa Fluor 633 anti-mouse (1:250, Invitrogen).

## QUANTIFICATION AND STATISTICAL ANALYSIS

### Data processing

For locomotion data, the forward, sideways, and rotational positions of the ball were computed in real time and acquired as voltage signals. Post hoc, these signals were converted to radians, unwrapped, low-pass filtered at 20 Hz, downsampled to 1,000 Hz, converted to degrees, and smoothed with a Gaussian window (σ = 200 ms). Directional velocities (Δposition/Δt) were calculated and smoothed with a Gaussian window (σ = 100 ms). Rotational velocity was then converted to °/s, and forward/sideways velocities to mm/s (based on the diameter of the spherical treadmill). The position of visual stimulus on the panorama, acquired in real time as a voltage signal, was converted to degrees post-hoc.

For electrophysiology data, the liquid junction potential was corrected post hoc by subtracting 13 mV from recorded voltages^91^. Cell firing rate was estimated by detecting spikes in the membrane voltage signal (findpeaks, MATLAB) and convolving the resulting binary spike train with either a Gaussian kernel (σ = 50 ms, window = ±500 ms) or a causal exponential kernel (τ = 125 ms, window = 0–1.25 s), each scaled such that the total area under the kernel was unity. A Gaussian kernel was used for all analyses except for the current injection experiments, where a causal kernel was used to better resolve the temporal relationship between manipulated neural activity and behavioral output.

### Classification of behavioral epochs

Movement epochs were identified using a Schmitt trigger applied to forward velocity. An epoch was defined to start when velocity rose above an upper threshold and to end when it fell below a lower threshold (0.1 mm/s). For behavioral recordings in intact flies, the upper threshold for running was 5 mm/s; for electrophysiology-behavior recordings, it was 3 mm/s; and for walking analyses, it was 1 mm/s. Running epochs shorter than 5 s were excluded. Stationary epochs were defined as all time points outside of running, including periods when forward velocity remained within ±0.25 mm/s to account for minor vibrations or residual ball motion.

During experiments in which the visual cue rotated in closed loop with the fly’s rotation, pursuit epochs were identified by calculating the mean object position within a 2-s sliding window (1.5-ms step size) and selecting periods when the mean position remained within ±35° of the midline while the fly was running^92,93^. Epochs with short gaps in pursuit (<2 s) were included. Epochs of pursuit shorter than 5 s were excluded, as were the first/last 2 s of each trial.

During experiments in which the visual object moved in open loop, the pursuit index^45^ was computed using a 5-s sliding window (50-ms step size) and defined as the product of “tracking fidelity” (the correlation between visual object position and the fly’s rotational velocity) and “vigor” (the net ipsilateral turning, normalized to 1). The first and last 5 s of each trial were excluded from analysis.

### Computing responses to open-loop visual stimuli

To assess behavioral and neural responses to open-loop visual stimuli, each animal’s average evoked response was calculated, segregated by relevant behavioral state (e.g., all, running only, or stationary only) during the visual stimulus presentation. Stimulus presentations were assigned to a behavioral state if >70% of the timepoints in the stimulus presentation met the criterion. To correct for an individual animal’s turning biases (e.g., a tendency to turn more left or right due to innate preference and/or placement on the ball), turn responses to left and right stimuli were mirrored and averaged. Visual response latency was defined as the time from the start of the visual motion pulse to the first peak in the time-derivative of firing rate, measured during stationary epochs at each cell’s preferred object position. For summary plots, each motion pulse was assigned to its center position (°): −123.75, −101.25, −78.75, −56.25, −33.75, −11.25, 11.25, 33.75, 56.25, 78.75, 101.25, 123.75, 146.25, where negative values represent positions in the left visual field. In some plots, to improve clarity, traces were slightly offset to the left or right of center along the x-axis to reduce overlap.

We computed the expected right–left firing rate difference (*R* − *L*) on a per-cell basis, assuming that right and left cells have mirror-symmetric tuning properties. In other words, for each fly, the response of the contralateral (mirror-image) cell was estimated by mirroring the visual stimulus. Specifically, if *R* and *L* denote the firing rates of cells in the right and left hemispheres, respectively, and these rates depend on object position (θ) and direction of object motion (rightward or leftward), then the assumption of mirror symmetry implies that

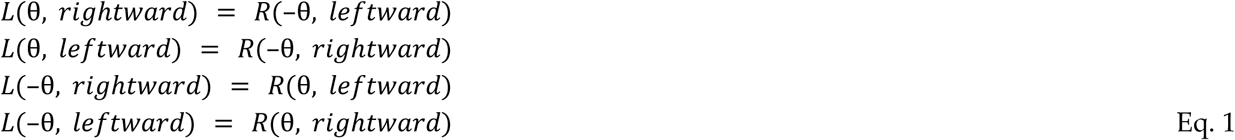

Positive and negative values of θ denote object positions in the right and left visual fields, respectively. Using these mirrored responses, we computed (*R* − *L*) for every motion-pulse stimulus in every fly. The associated uncertainty was determined by error propagation, where the propagated error equals the square root of the sum of squared standard errors from the individual components. In each fly, the right–left difference in AOTU019 firing rate (*R* − *L*) represents the contribution of AOTU019 to steering, and likewise, the right–left difference in AOTU025 firing rate (*R* − *L*) represents the contribution of AOTU025 to steering. This follows from Eq. 5 in our network model (see below).

The direction selectivity index (DSI) was calculated by comparing firing rate (FR) responses to progressive (away from the midline) versus regressive (toward the midline) motion at the same visual stimulus position^45^, expressed as the difference divided by the sum of the two responses per cell:

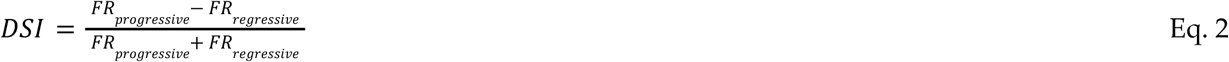

The signal-to-noise ratio (SNR) was computed on a per-cell basis. For each motion pulse sweep position, we measured the average evoked firing rate per trial during stationary epochs. Across trials, we then calculated the mean and variance of these per-trial averages. The SNR was defined as the ratio of the mean divided by the variance, and averaged across each cell’s preferred object positions.

### Computing responses to closed-loop visual stimuli

To standardize fixation angles across flies, visual object positions were centered at 0° by subtracting the mean visual object position during fixation. This procedure assumes that each fly is attempting to fix the object precisely at 0°, and so any small systematic biases in a fly’s mean fixation angle are due to imprecision in setting the fly’s azimuthal angle relative to the display screen, or other similar sources of measurement error. To analyze the relationship between visual object position and turning behavior during closed-loop fixation, average rotational velocity (adjusted for visuo-motor delay) was computed in 9° visual object position bins for each fly. Linear regressions were fit for visual object positions between ±20° by genotype, with slopes, confidence intervals, and Pearson correlation coefficients calculated per fly.

To quantify how tightly object positions were concentrated around the midline, we computed the median absolute deviation from 0° for each animal. Object positions during walking were pooled across time and trials for each condition (e.g., before versus during arousal stimulation). Here, smaller values indicate greater concentration of object positions near the midline.

### Computing the relationship between neural activity and behavior

To examine relationships between neural activity and behavior, average firing rates were computed for each fly in bins of 1 mm/s for forward velocity, 20°/s for rotational velocity, and 0.05 for the pursuit index to assess neural activity and pursuit performance. Similarly, average rotational velocity was computed for each fly in bins of 5 spikes/s. To assess temporal relationships between neural activity and behavior, we computed the cross-correlation between firing rate and lagged changes in directional velocity over a ±1 s window in 1.5 ms steps. For rotational velocity analyses, only ipsilateral (same-side) turns relative to the recorded neuron were considered. Cells were excluded from analysis of lag times if their cross-correlation did not exhibit a clear peak, defined as a minimum prominence of 0.15, because the lag associated with the peak correlation is not interpretable if the peak correlation is particularly weak. Data within 140 ms of walking onset or 250 ms before walking offset were excluded, as were the first and last 250 ms of optogenetic stimulation trials.

For current injection experiments, the average evoked response was calculated per fly for depolarizing and hyperpolarizing injections (1 s) within a 3 s window. Analyses were restricted to current injection trials during which the animal was actively walking (forward velocity >3 mm/s for more than 30% of the analysis window). Change in forward and rotational velocity was quantified by subtracting the average velocity during the 500 ms baseline period preceding current onset from the average during the final 500 ms of the pulse.

### Statistical analyses

Statistical analyses were performed in MATLAB. We used paired and two-sample Student’s t-tests, as well as ANOVAs to assess main effects, with post hoc Tukey-Kramer tests for pairwise comparisons where appropriate. For all repeated-measures ANOVAs, fly identity was modeled as a random effect using a linear mixed-effects model to account for within-animal variability. The p-values for all statistical tests are in **Table S1**, and are reported in the figures as: p < 0.05 (*), p < 0.01 (**), p < 0.001 (***), and p < 0.0001 (****).

### Data exclusion, sample sizes, and data reporting

For behavioral observations in intact animals, flies with <20 s of pursuit were excluded. For electrophysiology experiments, trials ran until the fly stopped walking or recording quality declined (10–120 min). Flies with <60 s of walking were excluded from analyses between neural activity and behavior. Cells showing minimal firing rate changes (<10 spikes/s) following depolarization or hyperpolarization were excluded from current injection analyses; individual current pulses in the lowest quartile of firing rate change (per cell) were also excluded.

Sample sizes followed field standards and pilot experiments that assessed variability and optimized stimulus and recording parameters. Reported means, medians, and s.e.m. values were computed across flies, not trials, after averaging each fly’s responses across stimulus repetitions. Replicates (n) thus indicate the number of flies, with a maximum of one neuron recorded per fly. The relatively large s.e.m. values for some visual responses (e.g., **Fig. 3D–F**) reflect broad variability across flies, likely due to differences in arousal-induced firing rate changes and behavioral engagement (some flies exhibited vigorous pursuit, while others walked minimally). Additional variability may arise from the aggressive dissections required for stable whole-cell recordings. All flies were included in these analyses.

### Analyzing connectome data

Cell type annotations, proofread reconstructions, synaptic connectivity, and neurotransmitter type predictions were taken from either the Flywire-FAFB^19–21,54,94–96^ (Female Adult Fly Brain) connectome (v783, https://ngl.flywire.ai/), the Flywire-BANC^24,97^ (Brain And Nerve Cord, female) connectome (v554), or the MaleCNS^23^ (Male Central Nervous System) connectome (v0.9, https://male-cns.janelia.org/). Data were analyzed using natverse packages^98^ (fafbseg v0.14.1, neuprintr v1.3.3, https://natverse.org/). Connections with <5 synapses were excluded from all analyses^22^. NeuronBridge^87^ was used to identify potential driver lines for targeting AOTU019 and AOTU025.

For each AOTU neuron postsynaptic to LC10a, we estimated the receptive field along the lobula’s medial–lateral axis by quantifying the distribution of its direct LC10a inputs. First, we defined a common retinotopic coordinate^25,83,99^ for the lobula by plotting all LC10a cells and mapping their full distribution as −15° to 165° (medial→lateral). For each AOTU cell, we visualized its presynaptic LC10a as a grayscale heatmap, with pixel intensity proportional to input synapse weight. The dorsal-most regions, which primarily contain cell bodies and axonal tracts, were excluded. The receptive field along the medial–lateral axis was computed as the per-column median grayscale intensity. Traces were then smoothed with a Gaussian filter.

Neuron cable width was analyzed by skeletonizing neurons from the Flywire-FAFB (v630) dataset using the “wave-front” skeletonization method^22^. Median primary dendrite widths were calculated for within-brain comparisons and visualized as normalized kernel density estimates. Measurements, based on dehydrated brain data, may underestimate true values.

### Network model

The model was run in MATLAB 2025a at 100 Hz, iterating 20,000 times per condition. The LC10a receptive field estimates for each AOTU cell were computed from male connectome data (**Fig. 3B, S8**), and were used to generate a lookup table of predicted neural activity as a function of the time-varying object position (θ(*t*)) in azimuthal space. Left and right copies of each AOTU cell were assumed to have mirror-symmetric visual receptive fields. The contribution of each AOTU cell was then weighted according to the number of synapses it forms onto DNa02 (averaged across hemispheres). Left and right copies of each AOTU cell were assigned identical output weights onto their ipsilateral copy of DNa02. We then modeled the activity of each DNa02 over time by linearly combining its AOTU inputs. Because AOTU019 is inhibitory and projects to the contralateral copy of DNa02, its contribution was sign-inverted and left-right inverted:

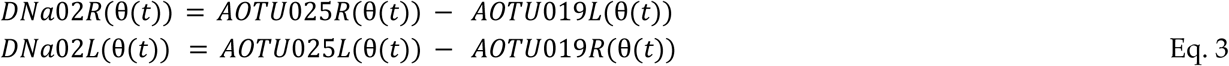

The fly’s rotational velocity command *r*(θ(*t*)) was taken as proportional to the right-left difference in DNa02 activity^48^:

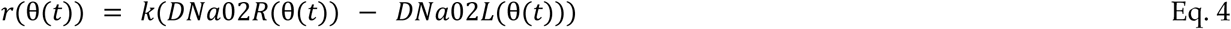

Note that combining Eq. 3 and Eq. 4 (and omitting (θ(*t*)) for clarity) yields

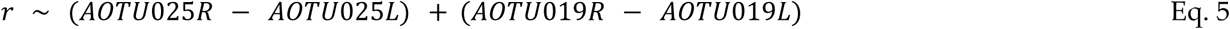

where the first quantity in parentheses represents the steering drive contributed by AOTU025, and the second represents the steering drive contributed by AOTU019. This is why we estimate each cell type’s contribution to steering drive by taking the right-left difference in the activity of that cell type (**Fig. 4D**).

In this model, *r*>0 indicates rightward (clockwise) steering. The scalar parameter *k* (0.45-0.65) defines the size of the rotational velocity response resulting from a given R-L difference in DNa02 activity. Rotational velocity was used to update the fly’s azimuthal orientation after a delay (Δt) of 200 ms, i.e., 20 steps of simulated time, to emulate the measured delay between neural activity and the rotation of the spherical treadmill. Rightward rotational velocity causes a negative (leftward) change in the object’s position θ(*t*), and vice versa. Thus, the object position evolves over time in response to the fly’s steering commands:

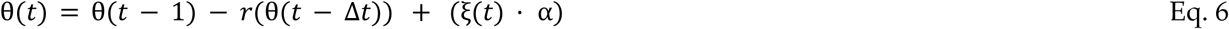

where ξ(*t*) is a random component scaled by a fixed scalar α. ξ(*t*) simulates the unpredictable movement of the visual target (e.g., a female fly during pursuit). For each run, ξ(*t*) was sampled from a normal distribution using the same frozen noise sample and then low-pass filtered at 0.05 Hz to generate a smoothed trajectory. Here, α controls the amplitude of perturbations to the object’s angular position. When α=0, the object remains fixed along the azimuthal axis unless displaced by the fly’s steering. As α increases, the object undergoes progressively larger angular shifts at each timestep. For simplicity, we quantified the mean object speed from its independent motion trajectory (i.e., without pursuer feedback) by averaging the absolute frame-to-frame change in object position over time, and report model runs according to this value.

To quantify the model’s ability to steer the object toward the midline and maintain this setpoint despite this perturbation, we calculated the probability of the object remaining within ±5° of the midline (0°). Additionally, to measure the relationship between visual object position and the steering output of the model, we calculated the average rotational velocity (accounting for visuo-motor delay) in 2° bins.

We also tested the model’s ability to correct varying displacements of the object. To eliminate the effect of the delay in the model, we began with holding the object at a fixed position (e.g., 100°) for 1 s before allowing a corrective response. In this simulation, ξ(*t*) was sampled from the same normal distribution as before, but the scale of the random component (α) was decreased 10-fold, to better illustrate the model dynamics resulting from overshooting. To quantify the model’s ability to restore a stable setpoint following a large perturbation, we calculated the settling time, defined as the time required for the object position in each run to stabilize within a ±2° tolerance and remain within this range for >1 s.

In the first model variant (**Fig. 5D-H**), we introduced direction selectivity to either the AOTU019 or AOTU025 pathways by systematically varying the direction selectivity index (DSI) from 0 to 1. For each neuron type, direction selectivity was implemented by applying a scaling penalty to the response to contraversive object motion, such that higher DSI values increasingly suppressed responses of the right-hemisphere cell to leftward object motion, and vice versa. For simulations where DSI was held constant, it was set to 0.2 (0.66 penalty to contraversive motion) based on electrophysiological measurements from AOTU019.

In the second model variant (**Fig. 7C**), we simulated the effect of selectively eliminating AOTU cells to mirror our genetic silencing experiments. Specifically, we tested the impact of removing AOTU019 alone, AOTU025 alone, both AOTU019 and AOTU025, or only the “minor” AOTU pathway. In these simulations, the “minor” pathway included all AOTU neurons postsynaptic to LC10a and presynaptic to DNa02, using each cell’s estimated visual receptive field derived from its LC10a inputs in the male connectome (**Fig. 3B, S7**), and scaling each cell’s activity by its output weight onto DNa02. As all “minor” inputs are predicted to be excitatory^54^, their contributions were summed to produce a combined AOTU drive to DNa02 from each hemisphere. Because AOTU002 projects to the contralateral copy of DNa02, its contribution was left-right inverted. For the right side (and omitting (θ(*t*)) for clarity):

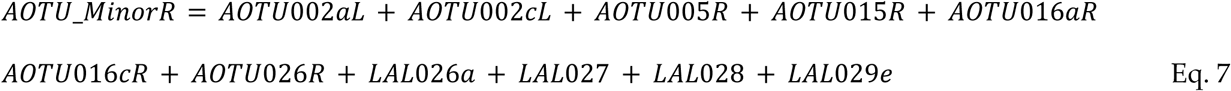

and equivalently for the left side.

## KEY RESOURCES TABLE

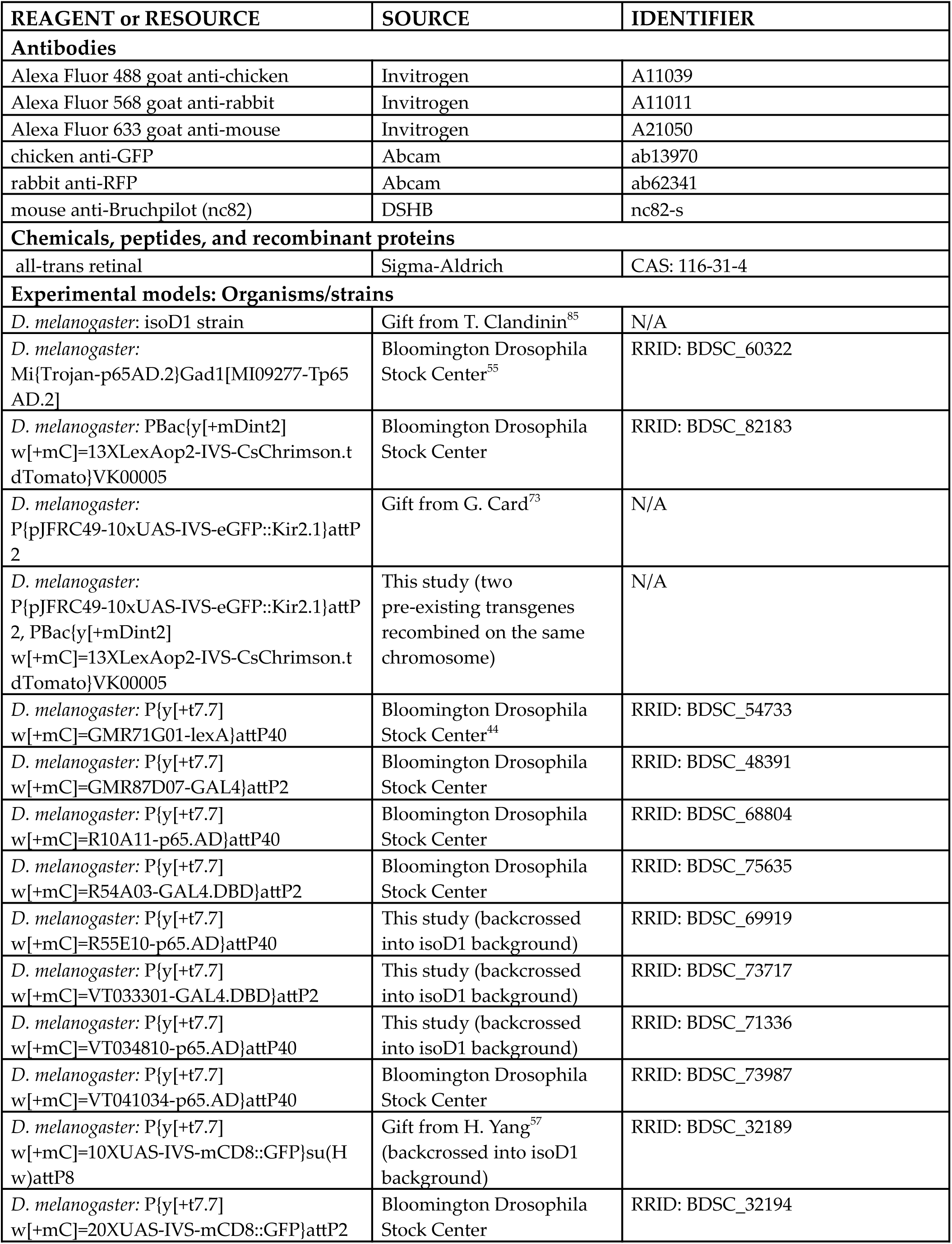
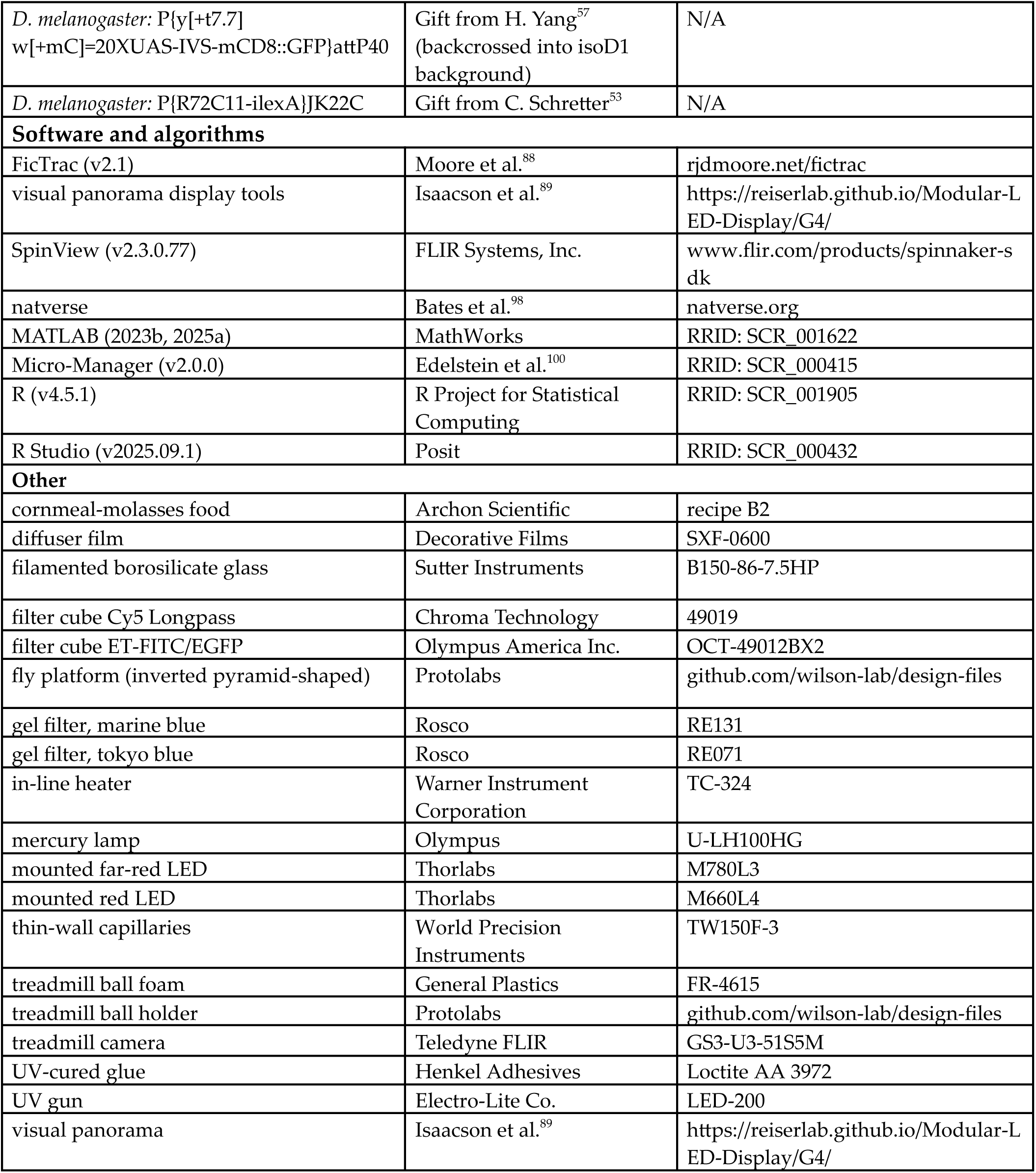

